# Haplotype Matching with GBWT for Pangenome Graphs

**DOI:** 10.1101/2025.02.03.634410

**Authors:** Ahsan Sanaullah, Seba Villalobos, Degui Zhi, Shaojie Zhang

## Abstract

Traditionally, variations from a linear reference genome were used to represent large sets of haplotypes compactly. In the linear reference genome based paradigm, the positional Burrows-Wheeler transform (PBWT) has traditionally been used to perform efficient haplotype matching. Pangenome graphs have recently been proposed as an alternative to linear reference genomes for representing the full spectrum of variations in the human genome. However, haplotype matches in pangenome graph based haplotype sets are not trivially generalizable from haplotype matches in the linear reference genome based haplotype sets. Work has been done to represent large sets of haplotypes as paths through a pangenome graph. The graph Burrows-Wheeler transform (GBWT) is one such work. The GBWT essentially stores the haplotype paths in a run length compressed BWT with compressed local alphabets. Although efficient in practice count and locate queries on the GBWT were provided by the original authors, the efficient haplotype matching capabilities of the PBWT have never been shown on the GBWT. In this paper, we formally define the notion of haplotype matches in pangenome graph-based haplotype sets by generalizing from haplotype matches in linear reference genome-based haplotype sets. We also describe the relationship between set maximal matches, long matches, locally maximal matches, and text maximal matches on the GBWT, PBWT, and the BWT. We provide algorithms for outputting some of these matches by applying the data structures of the r-index (introduced by Gagie et al.) to the GBWT. We show that these structures enable set maximal match and long match queries on the GBWT in almost linear time and in space close to linear in the number of runs in the GBWT. We also provide multiple versions of the query algorithms for different combinations of the available data structures. The long match query algorithms presented here even run on the BWT in the same time complexity as the GBWT due to their similarity.

## 1 Introduction

Traditionally, due to the similarity between human genomes, haplotypes have been represented using their differences with a linear reference genome. A haplotype’s deviations from a reference genome are represented as single nucleotide polymorphisms (SNPs) or short insertion and deletions (indels). These genetic variants are often represented as binary (or categorical) values at positions mapped to a linear reference genome. Pangenome graphs [25, 32], provide an alternative method of representing sets of genetic variations through the use of a graph. They have been proposed as an alternative to reference genomes that better captures the full-range of human diversity, possibly reducing reference bias [4]. In particular, the Human Pangenome project [68] aims to “create a more sophisticated and complete human reference genome with a graph-based, telomere-to-telomere representation of global genomic diversity.” A pangenome graph provides a natural way to encode insertions, deletions, copy number variations, inversions, and complex structural variants using a graph, as opposed to variations from a reference genome. Pangenome graphs have shown promising results when used in read mapping [32,39,57,63], genotyping structural variants and large indels [17,23,24,37,39,55, 63] (previously largely inaccessible variants), and motif search [67]. Construction [31,37,42,56] of pangenome graphs can be viewed as whole genome multiple alignment.

Recent advancements in DNA sequencing and genotyping technologies have made massive datasets of millions of human genomes increasingly available [36,64,65]. These large biobank scale datasets of genotyping and sequencing data are currently stored by their variations with respect to a linear reference genome. These datasets have many applications such as genealogical analysis [46, 70], imputation [59], phasing [12, 20, 43], Identity-by-Descent segment detection [14, 48, 64], and local ancestry [72]. These applications usually rely on efficient positional haplotype matching methods such as the Positional Burrows-Wheeler Transform [22].

An alternative method of storing haplotypes is representing them as paths through a pangenome graph. Storing haplotypes as paths through pangenome graphs instead of with their variations from a linear reference genome allows the representation of more complex variation. This includes large insertions and deletions, inversions, and repeats. Therefore, efficient haplotype matching methods on haplotype datasets represented using a pangenome graph are needed. However, haplotype matches in pangenome graph based haplotype sets are not well defined. In this paper, we define haplotype matches in haplotypes sets represented as paths through a pangenome graph by drawing from haplotype matches in linear reference genome based haplotype sets (the PBWT paradigm) and position agnostic haplotype matches (the BWT paradigm). We also provide algorithms for efficiently finding these matches in biobank scale haplotype datasets. These haplotype matching definitions and algorithms on haplotype sets represented with pangenome graphs allow the analysis of haplotype matches with structural variations.

Durbin’s Positional Burrows Wheeler Transform (PBWT) [22] represents a set of equal length binary strings. It allows efficient query of one vs. all and all vs. all set maximal and long positional matches, where a positional match is a continuous range at the same position on two haplotypes where the haplotypes have equal value [46, 61]. The PBWT has also been extended to larger alphabets [50]. To perform genealogical search on biobank-scale data, we require efficient algorithms for finding IBD segments. The PBWT is the basis for many such algorithms for sets of haplotypes represented by their variation from a linear reference genome [28, 47, 73].

There have been a couple of works for storing sets of haplotypes using a pangenome graph. The graph Burrows-Wheeler transform (GBWT) [62] is one such data structure, it represents each haplotype in the set as a path in the pangenome graph. The GBWT stores these paths reversed in a generalized FM index with a few modifications (where a generalized FM index of a set of strings is the FM index of their concatenation with end of string symbols at the end of each string). This allows fast locate, count, and extract queries of paths in the pangenome graph. The GBWT also compresses the alphabet at each node to the local alphabet. It is run length compressed and takes *O*(*r*) space, where *r* is the number of runs in the GBWT. However, computation of LF in the GBWT doesn’t have good time complexity bounds; the computation of *LF* from a node *v* takes time linear to the number of runs in node *v*.

Since the GBWT is a run length compressed BWT with compressed local alphabets, algorithms on a run length compressed BWT are relevant. In their r-index paper, Gagie et al. described data structures for exactly this, a Burrows-Wheeler Transform (BWT) in space proportional to the number of runs in the BWT [30]. The time complexities of count, locate, and extract in the r-index are almost linear. The r-index can count and locate all occurrences of a string in a linear amount of predecessor queries on a predecessor query of *r* elements with universe size *n*. They also provide a data structure in 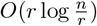 space that can extract *l* consecutive characters of the text in 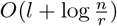 time. *n* is the size of the text of the BWT and *r* is the number of runs in it. In this paper, we apply the data structures of the r-index to the GBWT to allow efficient query in compressed space. The GBWT has generalized the PBWT’s compressed representation of large sets of haplotypes from the linear reference genome to a compressed representation of large sets of haplotypes as paths in a pangenome graph. However, the haplotype matching capabilities of the PBWT have not been replicated on the GBWT.

In general, algorithms for outputting matches to substrings of a query have been published for the BWT and the PBWT but not the GBWT. In the PBWT paradigm, Durbin defined set maximal matches, long matches, and locally maximal matches [22]. Algorithms for efficient set maximal and long match query were published later [46, 61]. Other substring match outputting algorithms in the PBWT paradigm include haplotype blocks [2, 49, 71] and k-SMEMs [10]. There are also PBWT algorithms for outputting haplotype blocks. Finally, the PBWT and many of its algorithms have been extended to compressed versions of the PBWT [9, 10, 18, 19, 69]. Related substring match outputting algorithms in the BWT include maximal exact matches (MEMs), maximal unique matches (MUMs), k-MEMs, k-rare MEMs, and long MEMs. These algorithms have also been extended to compressed representations of the BWT: MEMs [21, 41, 52, 53, 58], MUMs [33, 52], k-MEMs [52, 53, 66], k-rare MEMs [53], and only ouputting long MEMs [29].

In this work, we discuss new formulations of the maximal match types in the GBWT (and consequently BWT) paradigms. These are inspired from related matches in the PBWT paradigm. These new maximal match types result in generalizations of positional haplotype matching problems from the linear reference genome based PBWT to the pangenome graph based GBWT. We propose algorithms to efficiently solve them in sublinear space. In particular, we describe long and set maximal match query algorithms on the GBWT. We do this by extending the GBWT’s capabilities through the data structures of the r-index [30]. We use techniques similar to those of set maximal match queries on the BWT and PBWT and the long match query on the PBWT. We also note that the set maximal and long match query algorithms presented here can be straightforwardly modified to query a path in the GBWT vs. all other paths in the GBWT. Therefore, all vs. all set maximal match and long match queries can be performed in time close to linear to the sum of the lengths of the paths and the number of matches outputted (scaled by the predecessor query time). Furthermore, the set maximal match query we describe here doesn’t require the thresholds data introduced by Bannai et al. [5]. Finally, the long match query algorithm we present here on the GBWT applies to the BWT as well due to the structures’ similarity. It is the first efficient long match query algorithm on the BWT the authors are aware of.

## 2 Background

A *graph G* = (*V, E*) has a finite set of nodes *V* ⊂ ℤ and a set of edges *E* ⊆ *V* ^2^. We say a graph is *directed* if an edge (*u, v*) ∈ *E* can only be traversed in the forwards direction, *u* to *v*, and is not the same as the reverse edge, (*v, u*). A *path* in *G, P*, where *P* = *p*_0_*p*_1_ … *p*_*k*−*1*_ ∈*V* ^*k*^ and is of length *k*, is a sequence of nodes such that for each 0 ≤ *i* < *k* − 1, there exists an edge (*p*_*i*_, *p*_*i*+1_) ∈*E*. We use the notation *P* [*i*] = *p*_*i*_ to reference the *i*-th node on the path *P*. A *cycle* of length *k* is a path *C* of length *k* + 1 where *c*_0_ = *c*_*k*_, and a *directed acyclic graph* (DAG) is a directed graph which contains no cycles. A *pangenome graph* is a triple (*V, E*, seq), where (*V, E*) form a graph, and seq : *V* → `{A, C, G, T}`^∗^ is a function that maps each node to its associated genetic sequence. For a path *P*, we define seq (*P*) as seq (*P*) = seq (*P* [0]) seq (*P* [1]) … seq (*P* [*k* − 1]). That is, the sequence of a path in a pangenome graph is the concatenation of the sequences of each node the path visits. Figure 1 depicts a set of haplotypes, a possible pangenome graph representing this set of haplotypes, and a path in the pangenome graph that corresponds to each haplotype in the set. The rest of the paper assumes the reader is familiar with the Burrows-Wheeler transform (BWT) and haplotype matching problems on the positional Burrows-Wheeler transform (PBWT). For references, see [13, 22, 27, 61].

**Fig. 1.**
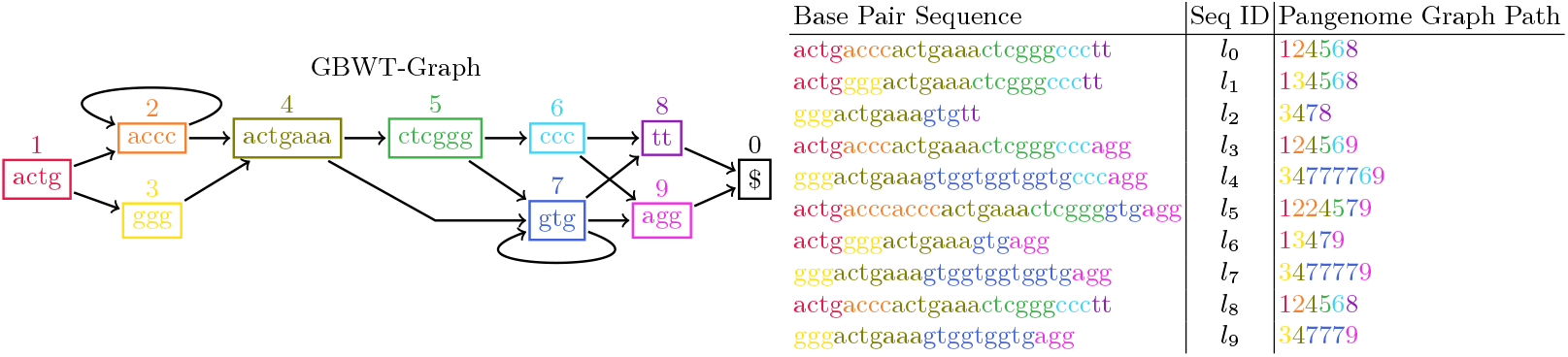
Haplotypes in a pangenome graph along with their sequence identifiers and the string representation of their path in the pangenome graph. Note that their may be many paths in a pangenome graph with equivalent sequences. The selection of a representative path for a haplotype is alignment of the haplotype to the pangenome graph.

### 2.1 GBWT

The GBWT represents a set of haplotypes as paths through a pangenome graph, it was introduced by Siren et al. [62]. It is well suited to represent sets of strings with high structural similarity and repetitiveness. It does this by representing strings as paths through a directed graph and storing only the paths and the graph. Nodes in the graph have string labels and the string corresponding to a path is the concatenation of the node labels. Finally, the GBWT is a run length compressed BWT of the reverse paths (where the paths are represented as strings over the alphabet of vertices in the graph). This run length compressed BWT is used to emulate a generalized FM-Index^1^. Since the GBWT represents strings as strings representing paths through a graph with nodes labeled by strings, the overloaded terminology results in hard-to-decipher sentences. Here we define terminology to make the formal writing of the paper more clear. We call the pangenome graph the GBWT represents the *GBWT-Graph*. We call the ultimate strings the GBWT is made to represent the *base-strings*, these are the whole genome haplotypes in a pangenome graph use case. Paths through the GBWT-Graph are *path-strings*, as they are both paths and strings in the emulated FM-Index. The emulated FM-Index of path-strings is the *GBWT-Index*. Finally, elements of the vertex set of GBWT-Graph and characters of path-strings are equivalent, therefore we refer to them as *node-characters*. As previously stated, node-characters are labeled with strings and the concatenation of the strings of the node-characters of a path-string is the base-string represented by the path-string. Finally, the GBWT is the GBWT-Index along with a representation of the base-strings, the GBWT-Graph is implicitly represented by the GBWT-Index (we say that an edge exists from node-characters *u* to *v* if and only if *uv* is a substring of a path-string). Note that the GBWT also may contain information on reversed node-characters, i.e., every node-character *v* has a corresponding reverse node-character 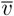. Furthermore, 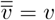. We call a GBWT where every path and its path-string reverse complement is contained in the GBWT a bidirectional GBWT. As Siren et al. note, a bidirectional GBWT is similar to an FMD-index as described by Heng Li [40]. A natural constraint for bidirectional GBWTs representing pangenome graphs is that the string of a node-character’s reverse should be the node-character’s string’s reverse complement.

Here we give a formal definition of the GBWT-Index. The GBWT-Index is built on a set of path-strings *S*_*P*_. It is a slight variation of a generalized FM index^2^ of the reverse of the path-strings in *S*_*P*_. The alphabet of the path-strings in *S*_*P*_ is the set of node-characters, *V*_*Σ*_. Furthermore, *V*_*Σ*_ = {0, …, *Σ*} and 0 is given the distinction of being the endmarker, it exists at the end of every path-string and only as the last character of a path-string. The base-string of the node-character 0 is $, the endmarker of the BWT. The main difference between the GBWT-Index and a regular multi-string BWT is that the GBWT-Index compresses the alphabet within runs in the *F* array. Note, the *F* array is the array of first node-characters in the sorted order of cyclic shifts of a text. For each node-character *v* ∈ *V*_*Σ*_, the range in the BWT of the run of *v* in the *F* array is referred to as *BWT*_*v*_. *BWT*_*v*_ is equivalent to *BWT* [*C*[*v*], *C*[*v* + 1]) for the classic FM-Index definition of the *C* array. Each *BWT*_*v*_ is stored separately and referred to as a *record*. Within each record, the alphabet of the *BWT* is compressed to the local alphabet. Define *Σ*_*v*_ as the set of node-characters in *BWT*_*v*_. Then, *f*_*v*_ is the local alphabet mapping. If *Σ*_*v*_ = |*Σ*_*v*_| is the number of unique node-characters in *BWT*_*v*_, then the local alphabet mapping, *f*_*v*_, is a monotone injective mapping from *Σ*_*v*_ to 0, …, *Σ*_*v*_ − 1 where *f*_*v*_(*w*) = *k* iff *w* is the (0-indexed) *k*-th smallest node-character in *Σ*_*v*_. Instead of storing *BWT*_*v*_ itself, it is mapped to its local alphabet. We call this mapped string 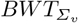 and 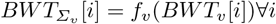 (and 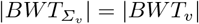). Then, 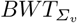 is stored in the *BWT*_*v*_ record run length compressed; every run of length *l* of characters *w* in 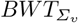 is stored as a pair (*w, l*). These runs are referred to as *BWT*_*v*_.*runs*. Lastly, each record *BWT*_*v*_ also stores *BWT*.*rank*(*v, w*) for all *w* ∈ *Σ*_*v*_. *BWT*.*rank*(*v, w*) is the total number of occurrences of *w* in *BWT*_*v*_ for all *v < v*. See Figure 2 for a depiction of a GBWT of the paths from Figure 1 and the relationship between the BWT and the GBWT.

**Fig. 2.**
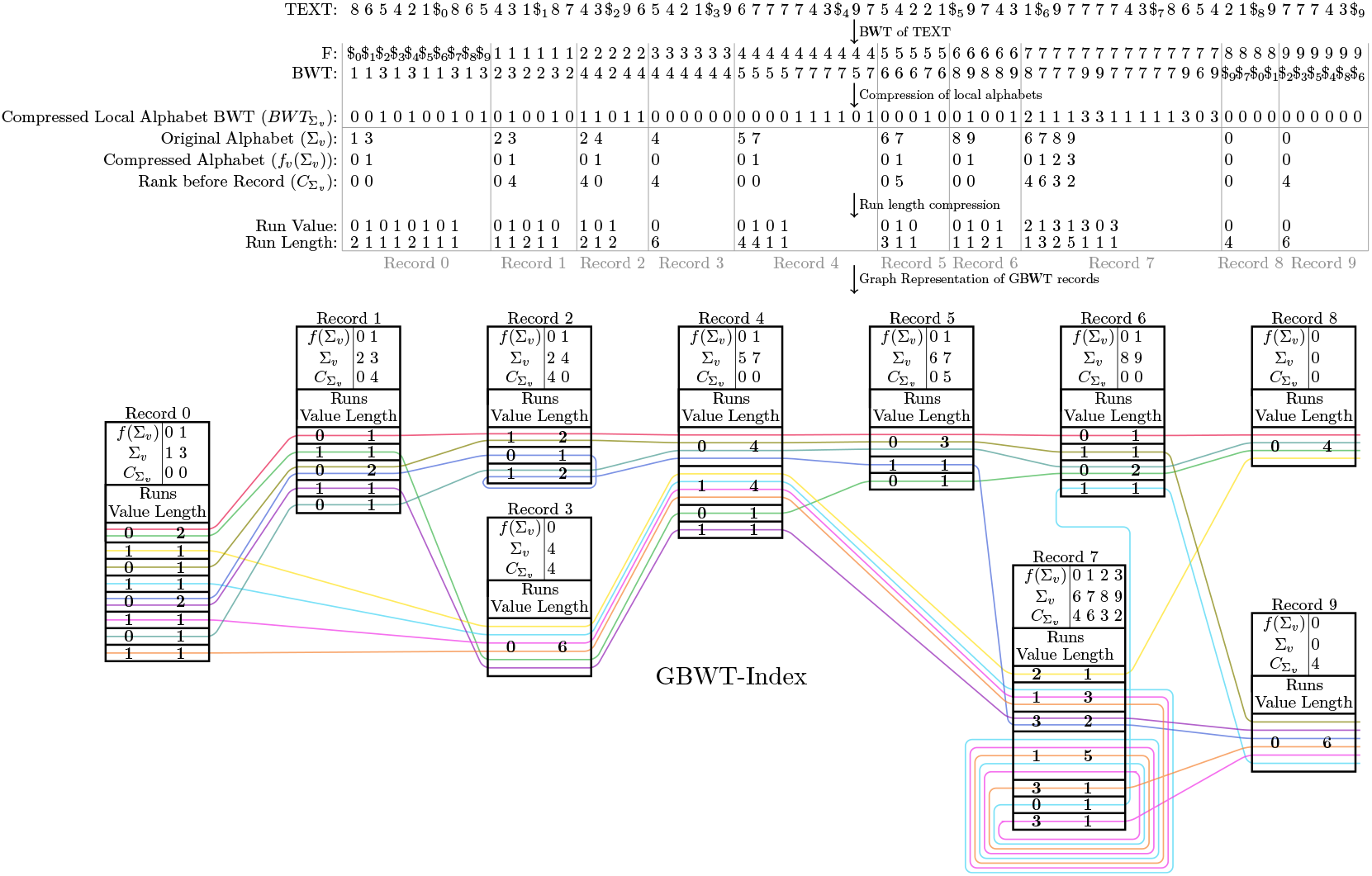
The relationship between the BWT and the GBWT. This figure depicts the GBWT of a set of paths through a pangenome graph. The underlying pangenome graph and their corresponding haplotypes are depicted in Figure 1. Note that in this figure, 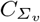 is used as shorthand to represent *BWT*.*rank*(*v*, _). The *i*-th value in 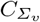 corresponds to *BWT*.*rank*(*v, w*) where *f*_*v*_ (*w*) = *i*.

In a bidirectional GBWT, the node-character set *V* is partitioned into two distinct parts, *V*_*f*_ and *V*_*b*_. *V*_*f*_ are the forward nodes and *V*_*b*_ are the backward nodes (in the original GBWT paper, backward nodes were referred to as reverse nodes, *V*_*r*_). *V*_*f*_ *⋃ V*_*b*_ = *V* and *V*_*f*_ *⋂ V*_*b*_ = ∅. Furthermore, for each node 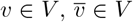 where 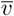 represents the complement of *v*. For each *v* and 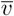 one is in *V*_*f*_ while the other is in *V*_*b*_. Lastly, 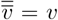 and 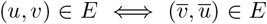. For any path-string *s* = *s*_0_*s*_1_.. *s*_*k*−1_, the reverse of *s* is rev(*s*) = *s*_*k*−1_*s*_*k*−2_.. *s*_0_ while the reverse complement is 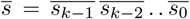. If *S*_*P*_ is the set of path-strings that a bidirectional GBWT is built upon, the set of strings in the GBWT-Index is 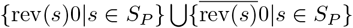. In other words, the generalized FM-Index is built on the set of strings of the reverse of every path-string and the reverse complement of the reverse of every path-string in *S*_*P*_ with the endmarker, 0, appended to the end of each.

From the values stored in the GBWT-Index, the *LF* mapping can be computed. As described, the *LF* mapping in the GBWT can not be computed without other data structures added to it. Positions in the GBWT are treated as node/offset pairs, (*v, i*) represents position *i* in *BWT*_*v*_; in the FM-Index, the equivalent position would be *C*[*v*] + *i*. Therefore, *BWT* [*C*[*v*] + *i*] = *BWT*_*v*_[*i*]. Then, *LF* ((*v, i*), *w*) = (*w, BWT*.*rank*(*v, w*) + *BWT*.*rank*((*v, i*), *w*)). If *w* ∈ *Σ*_*v*_ then *BWT*.*rank*(*v, w*) is stored in *BWT*_*v*_ and can be found by a predecessor query on *Σ*_*v*_, then the *BWT*.*rank*(*v, w*) can be computed directly. The *BWT*.*rank*((*v, i*), *w*) value is computed by iterating over the runs in *BWT*_*v*_. For all runs (*k, l*_*j*_) (call its start position *j*), in 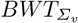, where *k* = *f*_*v*_(*w*), *BWT*.*rank*((*v, i*), *w*) = Σ_*j*|*j<i*_ min(*i, j* + *l*_*j*_) − *j*. This can be computed in *O*(*r*_*v*_) time where *r*_*v*_ is the number of runs in *BWT*_*v*_ by iterating through the runs. If *w* is not in *Σ*_*v*_, *BWT*.*rank*((*v, i*), *w*) = 0 and *BWT*.*rank*(*v, w*) can be computed in the following way: *BWT*.*rank*(*v, w*) = *BWT*.*rank*(*v, w*) where *v*′ = min {*j > v*|*w* ∈ *Σ*_*j*_}. *v*′ can be found by iterating through records after *v*, if no such node exists, then *BWT*.*rank*(*v*′, *w*) = *C*[*w* + 1] − *C*[*w*]. Note that in a bidirectional GBWT, *v*′ can be found by a successor query on 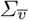 if the relative order of forward nodes and backward nodes are the same, 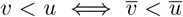. The time complexity of computing *LF* ((*v, i*), *w*) is *O*(*r*_*v*_) where *r*_*v*_ is the number of runs in *BWT*_*v*_ if *w* ∈ *Σ*_*v*_, otherwise it is *O*(|*V*|), therefore overall it is *O*(*r*_*v*_ + |*V*|) in the worst case (the time complexity is *O*(*r*_*v*_ + *t*_*s*_) where *t*_*s*_ is the time for successor query on 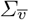 in bidirectional GBWTs), although in practice it runs much faster. Note that in many algorithms, the full LF is not necessary. For example, in count, locate, extract, and the set maximal match and long match queries presented here, the LF only needs to be computed if *w* ∈ *Σ*_*v*_. In these algorithms, the time complexity of LF can be said to be *O*(*r*_*v*_) in the GBWT.

### 2.2 Indexes on the GBWT from Previous Work

#### Document Array (DA)

The GBWT also stores a *document array DA*. The *DA* was described by Siren et al. in the original GBWT paper [62]. *DA*[(*v, i*)] returns the index of the path that *SA*[(*v, i*)] is a suffix of, i.e., if *SA*[(*v, i*)] points to a suffix of path *P*_*j*_, then *DA*[(*v, i*)] = *j*. Note that storing the complete document array would take *O*(*n*) space where *n* is the sum of lengths of paths in the panel. This is much larger than the rest of the GBWT due to the run length compression and high repetitiveness of genomic data, therefore Siren et al. sampled the document array only every *s* locations. For the 1000G and TOPMed datasets, values of *s* = 1, 024 and 16, 384 respectively resulted in document arrays of size roughly equal to the rest of the GBWT. The *DA* values are sampled at every *s* positions in the GBWT and the last position of every path.

#### FastLocate (FL)

Siren et al. modified the GBWT code after publication of the GBWT paper to add a new capability to the GBWT, FastLocate (FL). FL implements a modified version of the suffix array sampling done by the r-index. A high-level overview of the r-index data structure is that the suffix array is sampled at the beginning and ends of runs in the BWT. Then, a predecessor data structure on the sampled locations is stored. Each sampled location stores the suffix array value above it and the suffix array value below it. This allows the computation of the *ϕ* and *ϕ*^−1^ functions in the time needed to compute a predecessor. Note that unless otherwise noted, in this paper predecessor is inclusive, i.e., the largest value less than or equal to another. If *SA*[*k*] = *i*, then *ϕ*(*i*) = *SA*[*k* − 1] and *ϕ*^−1^(*i*) = *SA*[*k* + 1].

Note that any predecessor data structure on this data would yield the ability to compute *ϕ* and *ϕ*^−1^. For example, using Belazzougui and Navarro’s predecessor data structure from Theorem A.1 [7], a predecessor query can be answered in 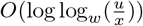 for a set of *x* integers in a universe of size *u* where a word takes *w* bits. This data structure takes *O*(*x*) words space. If this predecessor data structure is stored on the suffix array samples, *ϕ* and *ϕ*^−1^ can be computed in 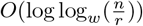 time in *O*(*r*) space. Navarro and Rojas-Ledesma published a useful reference on various solutions to the static and dynamic predecessor search problem [54].

Siren et al. only added the capability of computing the *ϕ*^−1^ function to the GBWT (the locateNext function). This takes roughly half the space of allowing both *ϕ* and *ϕ*^−1^ computation. This is done by storing a predecessor data structure on when suffixes are at the bottom of runs and sampling the values only at the top of runs. In other words, the theoretical suffix array of the GBWT is sampled at the beginning of every run in each *BWT*_*v*_. Note that, in their implementation, Siren et al. store (sequence id, offset) pairs instead of literal suffix identifiers, although these are functionally equivalent. Furthermore, the offset in a sample denotes the distance to the end of the path, i.e., (*i, j*) denotes the *j*-th suffix of rev(*P*_*j*_). The FastLocate implementation stores samples by GBWT runs instead of BWT runs. Call GBWT runs *o, o* ≤ *r* + |*V*_*Σ*_|, where *r* is the number of runs in the BWT of the text. |*V*_*Σ*_| ≤ *r*, therefore *r* and *o* are equivalent in asympotitic analysis and are used interchangeably. The *i*-th run (0-indexed) in the GBWT is the *j*-th (0-indexed) run of *BWT*_*k*_ where 0 ≤ *j < r*_*k*_ and 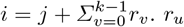 is the number of runs in *BWT*_*u*_.

The FastLocate structure is primarily composed of two structures: *samples* and *last. samples* is an array where *samples*[*i*] stores the (sequence id, offset) pair found at the beginning of the *i*-th run in the GBWT. *last* is the predecessor data structure on the universe of possible (sequence id, offset) pairs (in implementation, it is a sparse bit vector). Call the maximum length of a path in the GBWT plus one *l*_*max*_, then 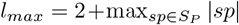. A pair (*i, j*) is encoded as *j* +*i*(*l*_*max*_). Therefore, *last* has length *n* = |*S*_*P*_ |*l*_*max*_. *last*[*k*] = 1 iff the pair 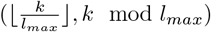 is at the bottom of a run in the GBWT. For all values *k* where *last*[*k*] = 1, the run id of the corresponding run is stored, we denote it as *last*[*k*].*runID*. Here we show how the *ϕ*^−1^ function is computed: given a sample (*i, j*), compute its (inclusive) predecessor, (*i, j*′), in *last*. Since (*i, k*) was never at the bottom of a run, the suffix below (*i, k* − 1) is (*a, b* − 1) if the suffix below (*i, k*) is (*a, b*) for *j*′ < *k* ≤ *j*. Therefore, the sample at the top of the run below *last*[*j*′ + *i*(*l*_*max*_)].*runID* is used. Precisely, 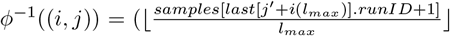, (*samples*[*last*[*j* + *i*(*l*_*max*_)].*runID* + 1] + *j j*) mod *l*_*max*_) where (*i, j*) is the inclusive predecessor of (*i, j*) in *last*.

### 2.3 Maximal Matches in the BWT and PBWT

In the BWT paradigm, there is a pattern string, *P*, and a text *T*. The BWT is built on the text and the pattern is queried against the text. In the PBWT paradigm, there is a query haplotype, *x* and a set of haplotypes *X*, all haplotypes are binary strings of equal length. The query haplotype is queried against the haplotypes in *X*. In the BWT paradigm, there are maximal exact matches, while in the PBWT paradigm, there are set maximal matches, long matches, and locally maximal matches. In the PBWT paradigm, matches must have the same position in the strings that they occur in, while in the BWT paradigm, matches are independent of the position they occur. Other than the difference on positional requirements, set maximal matches in the PBWT and maximal exact matches in the BWT are equivalent. However, there exist no analogous matches in the BWT for long matches and locally maximal matches in the PBWT.

A maximal exact match (MEM) is a match between the pattern *P* and the text *T* such that there is no larger substring in *P* that encompasses it and exists in *T*. Formally, a match can be defined as a triple (*i, j, k*) where *i* is the starting position of the match in *P, j* is the starting position of the match in *T, k* is the length of a match, and *P* [*i, i* + *k*) = *T* [*j, j* + *k*). Then, a match (*i, j, k*) between *P* and *T* is a MEM iff *P* [*i* − 1, *i* + *k*) does not exist in *T* and *P* [*i, i* + *k* + 1) does not exist in *T*. In the PBWT paradigm, a set maximal match between the query haplotype *x* and the set of haplotypes *X* is a match between *x* and *x*_*i*_ ∈ *X* s.t. no larger substring of *x* that contains this match occurs in *X* at the same position. A match in the PBWT between two haplotypes is only defined on starting position and length, (*i, k*), since they must occur at the same position in the haplotypes. Therefore, a match (*i, k*), between *x* and *x*_*i*_ ∈ *X* is a set maximal match between *x* and *X* iff ∀*h* ∈ *X, h*[*i* − 1, *i* + *k*) ≠ *x*[*i* − 1, *i* + *k*) and *h*[*i, i* + *k* + 1) ≠ *x*[*i, i* + *k* + 1). Therefore the idea behind set maximal matches in the PBWT and MEMs in the BWT are matches that can’t be extended in the query.

Another way to define matches are matches that can’t be extended in the query and matched location simultaneously. These are locally maximal matches in the PBWT. A locally maximal match, (*i, k*), between haplotypes *a* and *b* is a match that can’t be extended between these haplotypes. *a*[*i* − 1, *i* + *k*) ≠ *b*[*i* − 1, *i* + *k*) and *a*[*i, i*+*k*+1) ≠ *b*[*i, i*+*k*+1). An analogous match may be defined in the BWT paradigm. A locally maximal exact match (LEM) between a pattern *P* and text *T* is a match (*i, j, k*) s.t. *P* [*i* − 1, *i* + *k*) ≠ *T* [*i* − 1, *i* + *k*) and *P* [*i, i* + *k* + 1) ≠ *T* [*i, i* + *k* + 1). A LEM is a match such that no larger match encompasses it with the same relative position between pattern and text.

In the PBWT, long matches are defined. Long matches are locally maximal matches at least length *L*. Long matches are a natural filtering of locally maximal matches as there are typically many very small and irrelevant locally maximal matches. We call long matches in the BWT paradigm long LEMs. A long LEM is a LEM of at least length *L*. Note that Gagie recently published a paper on an algorithm for only outputting long MEMs while ignoring short ones [29]. This is not the same as outputting long LEMs as there may be many more long LEMs than long MEMs (but every long MEM is a long LEM).

Finally, matches that can’t be extended in the text may be of interest. We refer to these as a text maximal match. In the PBWT, text maximal matches are equivalent to locally maximal matches due to the restriction of matches occurring in the same position in the query haplotype and matched haplotype. However, in the BWT, text maximal matches may be different from LEMs. We refer to text maximal matches in the BWT paradigm as text maximal exact matches (TEM). A TEM between pattern *P* and text *T* is a match (*i, j, k*) s.t. no larger match encompasses it in the text, i.e., *T* [*j* − 1, *j* + *k*) does not exist in *P* and *T* [*j, j* + *k* + 1) does not exist in *P*. Note that TEMs and MEMs are inverses of one another. The MEMs of a pattern *P* with respect to text *T* are the TEMs of the pattern *T* with respect to the text *P*. Lastly, MEMs and TEMs are both LEMs but there maybe LEMs that are neither TEM nor MEM.

## 3 Methods

### 3.1 Haplotype Matching in Graphs

Here, we generalize the definitions of matches in the BWT and PBWT paradigms to the GBWT. We show that the GBWT in fact can mirror most, if not all, of the strengths of the PBWT, by solving the following four generalizations of the PBWT haplotype matching problems: one vs. all set-maximal and long matches and all vs. all set-maximal and long matches. For a GBWT with a panel of path-strings *P*, the one vs. all algorithms outputs all matches between a query path-string *Q* and *P* while the all vs. all algorithms output matches between *P*_*i*_ and *P* \ *P*_*i*_ for all *P*_*i*_ ∈ *P*.

Set-maximal and long path-string matches in GBWT are defined below. Note that if one views path-strings only as strings, these definitions are equivalent to that of MEMs and long LEMs in the BWT paradigm respectively. Similarly, if one views path-strings only as paths, these definitions are equivalent to that of set maximal matches and long matches in the PBWT paradigm. However, in path-strings, node-characters act as both characters and nodes in the underlying pangenome graph. Therefore, node-characters serve as a pseudoposition in the genome, if two haplotypes have the same node-characters in different literal positions in their genome, the equivalent node-character implies the position of these path-strings are equivalent in the underlying graph (which may have some underlying genomic meaning). Using these properties, a path-string match is a middle ground between the positionless BWT paradigm, and the absolute equivalent position PBWT paradigm. A path-string match is positionless in the strings, and an absolute position match in the graph. Therefore, when referring to path-string matches in the GBWT paradigm, we use locally maximal matches, long matches, and set maximal matches (the PBWT paradigm match terms) interchangeably with LEMs, long LEMs, and MEMs (the corresponding BWT paradigm match terms respectively).

#### Path-String Matches

A *match* between path-strings *A* and *B* is a triple of integers, (*i, j, k*), where 0 ≤ *i < i* + *k* ≤ |*A*|, 0 ≤ *j < j* + *k* ≤ |*B*|, and *A*[*i, i* + *k*) = *B*[*j, j* + *k*). In this 3-tuple, *i* represents the index of the first node-character in the path-string match in *A*, while *j* represents the index of the first node-character in the path-string match in *B*, and *k* represents the number of node-characters in the path-string match. A path-string match is *locally maximal* if it can’t be extended in either direction, i.e., (*i* = 0 ∨ *j* = 0 ∨ *A*[*i* − 1] ≠ *B*[*j* − 1]) ∧ (*i* + *k* = |*A*| ∨ *j* + *k* = |*B*| ∨ *A*[*i* + *k*] ≠ *B*[*j* + *k*]). Thus, there is no larger path-string match between *A* and *B* that fully contains *A*[*i*.. *i* + *k*) and *B*[*j*.. *j* + *k*) and starts in the same relative location.

Given a minimum length *L*, a *long match* is a locally maximal path-string match ((*i, j, k*) between *A* and *B*) where the length of the match is at least *L*. Long matches may be defined on other notions of length of a match. A natural one in the GBWT is the length of the underlying base-string. Then, for a node-character *v*, define its weight *w*_*v*_ as the length of *v*’s sequence, |seq (*v*)|. A match is long if the weight of the matched nodes in the match sum to at least *L*, that is, 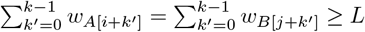.

When comparing a set of path-strings *P* = {*P*_0,_ *P*_1_,…*P*_*m-*1_} with a single path-string *Q*, we say a match (*i, j, k*) between *Q* and *P*_*x*_ is *set-maximal* if there does not exist a match (*i*′, *j*′, *k*′) between *Q* and *any P*_*y*_ ∈*P* such that [*i*.. *i* + *k*) ⊂ [*i*′.. *i*′ + *k*′). Note that this implies set-maximal matches are locally maximal. In the all vs. all matching problems, when considering set-maximal path-string matches for a path-string *P*_*x*_ ∈ *P* in the panel, the set of path-strings being compared against is all other path-strings in the panel, *P* \ {*P*_*x*_}.

### 3.2 Set Maximal Match Query and Supporting Indexes

Outputting MEMs in the BWT is frequently done through the computation of matching statistics [5, 58]. Due to the similarity between the GBWT and the BWT, these techniques apply to the computation of set maximal matches and matching statistics in the GBWT as well. In this paper, we compute set maximal matches through the computation of virtual insertion positions and their corresponding LCP values. These are closely related to matching statistics, from matching statistics it is straightforward to get virtual insertion positions and their LCP values and vice versa. However, the method of computing virtual insertion positions and their LCP values presented here does not use thresholds. Thresholds are auxiliary data that take *O*(*r*) space introduced by Bannai et al. to enable the computation of matching statistics in the BWT [5]. Our method efficiently computes matching statistics without thresholds. The method of computing set maximal matches in the GBWT we present relies on four main capabilities: mapping in the GBWT, the suffix array, the LCP array, and access to the text. For mapping in the GBWT, we require inverse LF and generalized LF computation. We describe Fast LF (FLF), a structure that provides these capabilities with time complexities better than the GBWT. We also present Fast LCP (FLCP), an augmentation to the GBWT that allows efficient computation of the *ϕ* and *LCP* functions. Lastly, we apply a compressed text (CT) random access data structure designed for the BWT [30] to the GBWT. The inverse LF capability is only required if text access is not available. See Table 1 for a summary of these structures and the functions the provide.

**Table 1.**
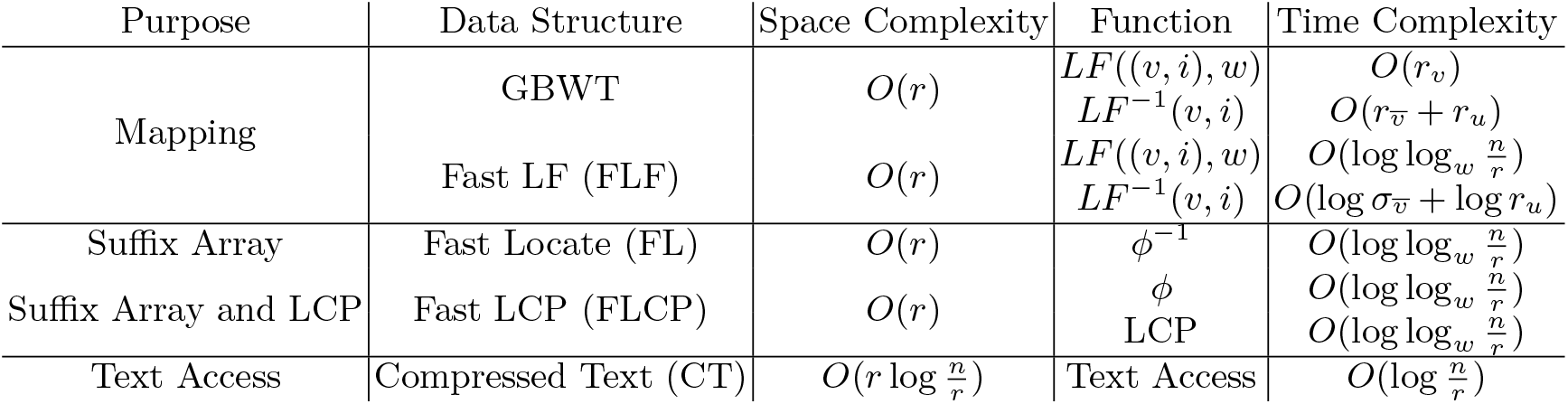
Data structures for virtual insertion positions and LCP values. Various data structures used in set maximal and long match queries along with their space complexities, purposes, functions, and corresponding time complexities.

| Purpose | Data Structure | Space Complexity | Function | Time Complexity |
| --- | --- | --- | --- | --- |
| Mapping | GBWT | $O(r)$ | $LF((v, i), w)$ | $O(r_v)$ |
| | | | $LF^{-1}(v, i)$ | $O(r_{\bar{v}} + r_u)$ |
| | Fast LF (FLF) | $O(r)$ | $LF((v, i), w)$ | $O(\log \log_w \frac{n}{r})$ |
| | | | $LF^{-1}(v, i)$ | $O(\log \sigma_{\bar{v}} + \log r_u)$ |
| Suffix Array | Fast Locate (FL) | $O(r)$ | $\phi^{-1}$ | $O(\log \log_w \frac{n}{r})$ |
| Suffix Array and LCP | Fast LCP (FLCP) | $O(r)$ | $\phi$ | $O(\log \log_w \frac{n}{r})$ |
| | | | LCP | $O(\log \log_w \frac{n}{r})$ |
| Text Access | Compressed Text (CT) | $O(r \log \frac{n}{r})$ | Text Access | $O(\log \frac{n}{r})$ |

We present five versions of set maximal match query on the GBWT. All set maximal match algorithms can be understood with Algorithm 1 using various versions of the virtual insertion, lcp, and output matches algorithms. The first operates on the GBWT alone. It calls Algorithm 2 and provides it with a GBWT to compute the LF function. Then it calls Algorithm 3 and provides it with a GBWT (bidirectional) to compute the inverse LF function. Lastly, it calls Algorithm 4 and provides it a DA to perform the locate algorithm. The second runs on the GBWT as recently modified by Siren et al. This adds the FastLocate functionality which stores samples at the beginning of runs in the GBWT. This is done by providing the OutputMatches function with both the GBWT and the FL. The third version modifies the FastLocate functionality to store samples at the beginning and ends of runs as originally described by Gagie et al. [30]. We also add samples of the longest common prefix array at the beginning of runs in the GBWT. This modification is referred to as Fast LCP (FLCP). In this version, the Virtual insertion algorithm is called with Algorithm 5 and also provided the FastLocate data structure to return the suffix values at the virtual insertion positions. Then, the matches are outputted with Algorithm 6, which makes use of the FLCP. These versions of set maximal match query don’t have good time complexity bounds since they must compute LF by iterating through runs within records in the GBWT. Therefore, the fourth version augments the GBWT through the use of a predecessor data structure to enable more efficient LF computation. This modification of the GBWT is referred to as Fast LF (FLF). The last version uses access to the text to obtain a runtime closer to linear. This is done by computing the LCP values using a compressed data structure that allows efficient random access to the text (see Algorithm 7). The compressed text data structure (referred to as CT here) was introduced by Gagie et al. [30]. The first four versions require a bidirectional GBWT as input while the last will run on any GBWT.

#### Algorithm 1

Set Maximal Match Query: Outputs all set maximal matches of a query path *Q* and a GBWT

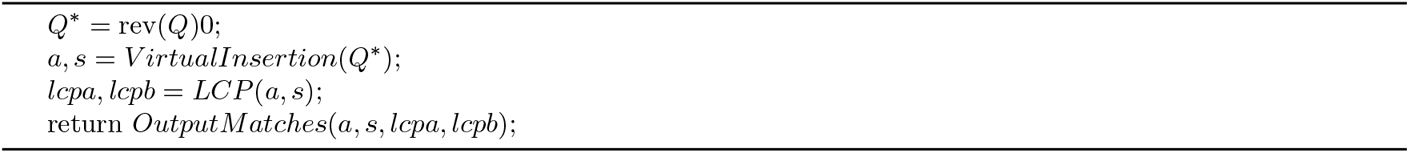

#### GBWT + DA: Set Maximal Match Query using only the GBWT and DA

Here we discuss the first version of the set maximal match query. It has two high level steps. Note that the traditional BWT sorts suffixes of the original strings. The GBWT is built on the reverse of the original path-strings and therefore it sorts the reversed prefixes of the original text (the PBWT sorts in this way as well [22]). In the first step, the query path-string is “virtually inserted” into the GBWT. For each reversed prefix of the query path-string, virtual positions are computed. The virtual position of a reversed prefix is a position in the GBWT where the longest common suffix with the suffix at or above the virtual insertion position is it’s longest common suffix with all other reversed prefixes in the GBWT. In the second step, the longest common prefix of each reversed prefix of the query and the adjacent reversed prefixes in the GBWT is computed. When the length of the longest common prefix of reversed prefix *i* is greater than or equal to the length of longest common prefix of reversed prefix *i* + 1, there is a set maximal match ending at position *i*.

The first step is to “virtually insert” the query path-string, *Q* into the GBWT. Before doing so, the query path-string is reversed, and then has an endmarker 0 appended to it (call this new path-string *Q*^∗^), as has been done to all other path-strings in the GBWT before construction. We define suffix *k* of *S* as *S*[*k*.. |*S*| − 1] and reversed prefix *k* of *S* as *S*[*k*.. 0]. Then suffix |*Q*^∗^| − 1 − *k* of *Q*^∗^ is equivalent to reversed prefix *k* of $*Q* for 0 ≤ *k <* |*Q*^∗^|. Then, we obtain, for all 0 ≤ *k <* |*Q*^∗^|, a position that the longest common prefix of suffix *k* of *Q*^∗^ could sort to in the GBWT if it contained it. Call *a*_*k*_ = (*v, i*) the position in the GBWT that the longest common prefix of suffix *k* of |*Q*^∗^| sorts to in the GBWT (the *i*-th position in node *v*). Then, *a*_*k*_−_1_ can be computed directly from *a*_*k*_ with the generalized LF function, *a*_*k*_ −_1_ = *LF* ((*v, i*), *Q*^∗^[*k* − 1]) if *Q*^∗^[*k* − 1] ∈ *Σ*_*v*_, (*Q*^∗^[*k* − 1], 0). Otherwise, we set *a* _| *Q∗* | 1_ = (0, 0), the first position in the endmarker node. Therefore, if the generalized LF function can be computed in *O*(*t*_*LF*_) time, the *Q*^∗^ string can be “virtually inserted” into the GBWT-Index in *O*(|*Q*^∗^| *t*_*LF*_) time. See Algorithm 2 for pseudocode of the virtual insertion algorithm with the GBWT.

##### Algorithm 2

Virtual Insertion: Given string *t* and a GBWT or FLF, return array *a* where *a*[*i*] is the virtual insertion position of *t*[*i* …, |*t*| − 1]

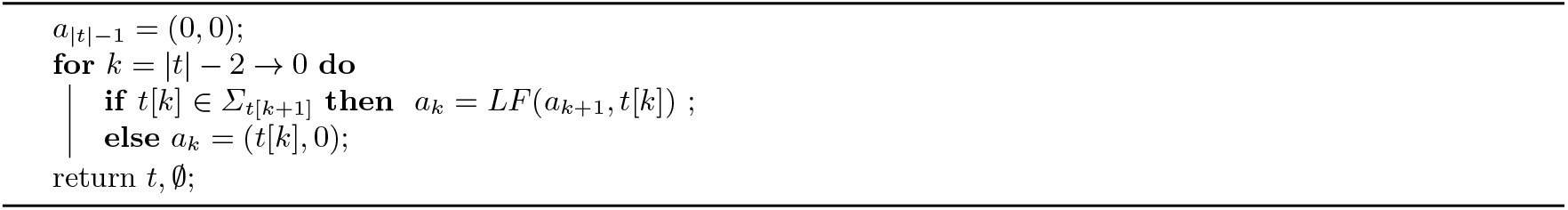

Now, for each suffix *k* of *Q*^∗^, we compute the values *lcpa*_*k*_ and *lcpb*_*k*_. If *a*_*k*_ = (*v, i*), *lcpa*_*k*_ is the length of the longest common prefix of suffix *k* of *Q*^∗^ and the suffix at position (*v, i* − 1). *lcpb*_*k*_ is the length of the longest common prefix of suffix *k* of *Q*^∗^ and the suffix at position (*v, i*). *lcpa* and *lcpb* represent longest common prefix above and below respectively. Note that *lcpa*_*k*_ ≥ *lcpa*_*k*−1_ − 1 (and *lcpb*_*k*_ ≥ *lcpb*_*k*−1_ − 1). Therefore, we iterate *k* from 0 → |*Q*^∗^| − 1 computing *lcpa*_*k*_ and *lcpb*_*k*_. In order to do this, we use the inverse LF function, *LF* ^−1^, to recover the text of the suffixes at positions (*v, i* − 1) and (*v, i*). This is where the bidirectional requirement for the input GBWT comes from, bidirectional GBWTs allow efficient inverse LF computation. See Algorithm 3 for pseudocde of LCP computation in a bidirectional GBWT.

Finally, when max(*lcpa*_*k*_, *lcpb*_*k*_) ≥ max(*lcpa*_*k*_−_1_, *lcpb*_*k*_−_1_), there is a set maximal match to output with *Q*^∗^[*k*.. *k* + max(*lcpa*_*k*_, *lcpb*_*k*_) − 1]. Then, the suffixes in the GBWT that contain this path-string are found in the typical BWT search method, repeatedly LFing over the substring. I.e., all the set maximal matches are outputted and found by a locate query on *Q*^∗^[*k*.. *k* + max(*lcpa*_*k*_, *lcpb*_*k*_) − 1]. Note that the GBWT and DA do not support outputting the location of matches within paths, therefore this version of the set maximal match query version does not output the location of matches within the paths that match the query path. This would be remedied if document array samples were replaced with suffix array samples. See Algorithm 4 for pseudocode for outputting matches using the GBWT given virtual insertion positions and their corresponding computed LCP values.

There are three main components of the set maximal match query algorithm. First is the virtual insertion, where positions of the reversed prefix of *Q* sort in the GBWT. Second is the computation of LCP values of *Q* and below *Q* for all reversed prefixes of *Q*. Lastly, there is the outputting of matches to paths in the GBWT, set maximal matches are outputted for paths that match *Q* on the set maximal match range. Note that the text identifier samples the DA contains do not contain information about the location of the sample within the path, therefore set maximal matches outputted by this version of the query only output the starting location in the query, the length of the set maximal match, and the haplotype that contains the match (this 3-tuple is outputted once for every match that exists in this haplotype). The virtual insertion takes *Q* LF operations. LCP computations may takes *O*(|*Q*|^2^) inverse LF operations in the worst case. The find operation takes LF operations proportional to the sum of the lengths of the ranges of set maximal matches. Lastly, the locate operation takes *O*(*s*) operations per match outputted. where the document array is sampled at every *s* positions. Therefore, this version of the set maximal match query algorithm takes 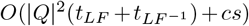 where *t*_*LF*_ and 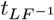 are times taken for LF and inverse LF computation respectively and *c* is the number of matches outputted. Note that *t*_*LF*_ and 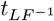 are *O*(*r*_*v*_) and 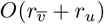 respectively, where *v* is the GBWT node that the operation was run from and *u* is the node in the result of the inverse LF computation.

##### Algorithm 3

LCP: Given Virtual Insertion positions *a* of string *t*, compute LCPs using *LF* ^−1^

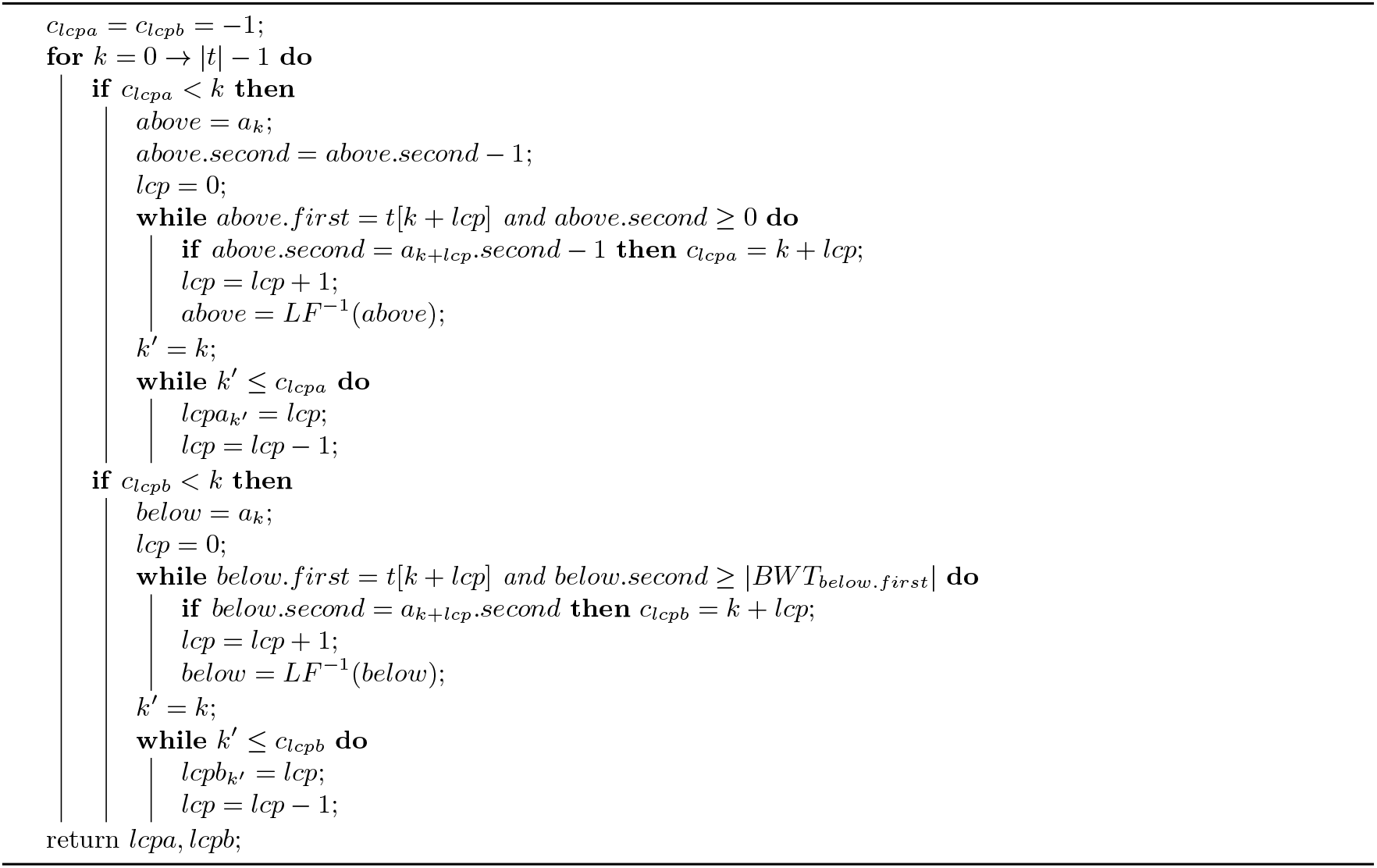

##### Algorithm 4

OutputMatches: Output Matches using a GBWT or a GBWT and FastLocate given virtual insertion positions *a* and LCP values *lcpa, lcpb*

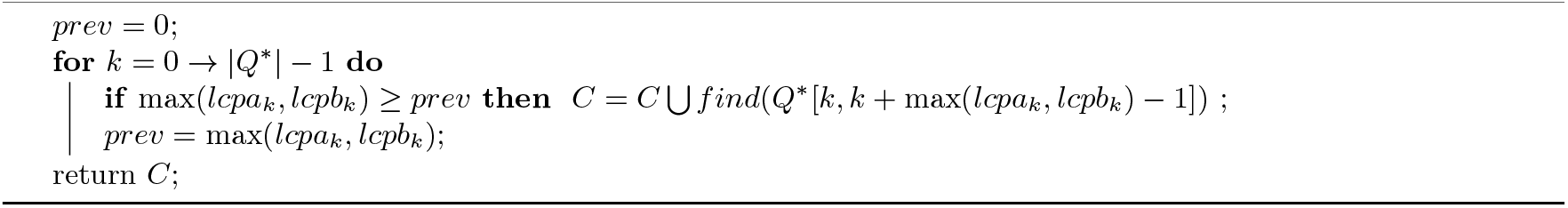

#### GBWT + FL: Replacing the Document Array with FastLocate

The second version of the set maximal match query algorithm makes use of the FastLocate functionality added to the GBWT after publication. The only changes to the original algorithm this makes are in the final part of the algorithm, when outputting found set maximal matches. This also requires keeping track of the suffix when finding all sequences that contain a set maximal match. See Algorithm 5 for pseudocode for keeping track of the suffix corresponding to virtual insertion positions. This only affects the find and locate queries computed on the GBWT at the end. The find query now needs to keep track of the suffix array value of the first position in the found range. This can be done easily while computing the *LF* function during extension of the match without changing its time complexity. Since the FastLocate structure stores suffix array values, it stores information about where on the path a prefix starts. This allows this version of the set maximal match query algorithm to output the starting positions of matches on the paths the query path matches with. Therefore, the use of the FastLocate data structure removes the need for the document array sampling and replaces the previous *O*(*cs*) time complexity for outputting matches with locate to (*c*(*tp* + |*BWT*0|)) (the + *BWT*0 is because in a bad case it may be necessary to iterate through large runs in the endmarker to find the start position for a match due to the way FastLocate stores positions). It also changes the space complexity from 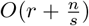 to *O*(*r*). Note that in practice, the FastLocate structure is a few times larger than the sampled document array for 1000G data. Furthermore, 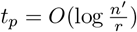 in the GBWT implementation and the bad case for locate almost never happens in practice. Here *n* = *Σsp S sp* + 1 is the length of the text and *n*′ = |*SP*|∗ *lmax* is the length of the bit vector for the predecessor data structure in FastLocate.

##### Algorithm 5

Virtual Insertion: Given string *t* and a FastLocate and a (GBWT or FLF), return arrays *a* and *s* where *a*[*i*] is the virtual insertion position of *t*[*i* …, |*s*|− 1] and *s*[*i*] is the suffix value at that position

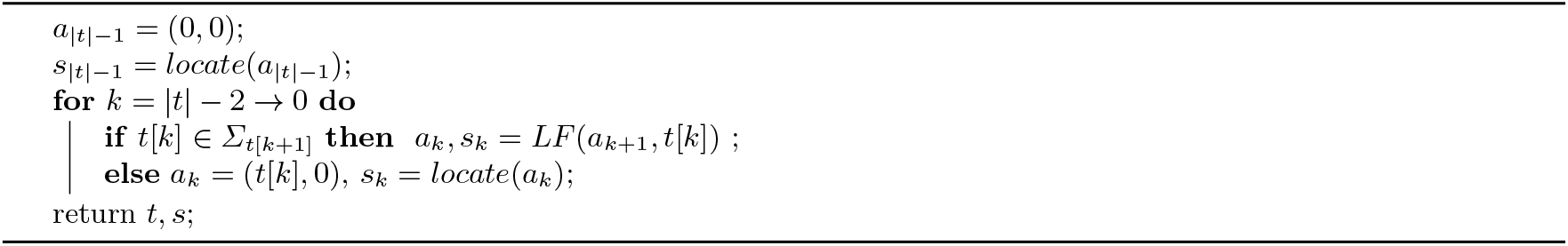

#### GBWT + FL + FLCP: Adding Fast LCP

The third version of the set maximal match query algorithm modifies the FastLocate functionality. It adds samples of suffixes found at the bottom of the runs, *samplesbot*, and a predecessor data structure on the suffixes that occur at the top of runs, *first*. These definitions are similar to *samples* and *last* from FastLocate, see Section 2.2. *samplesbot*[*i*] is the suffix array value at the bottom of the *i*-th run in the GBWT. *first* is a sparse bit vector where 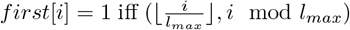 is at the top of a run (or equivalently, contained in *samples*), where *l*_*max*_ = 2 + max_*sp*∈*SP*_ |*sp*|. These data structures are added to allow the computation of the *ϕ* function. They maintain the space complexity of the FastLocate data structure, but roughly double its size. The *ϕ* function can be computed similarly to the *ϕ*^−1^ function, 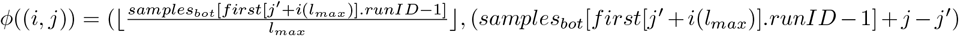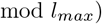,where (*i, j*) is the predecessor of (*i, j*) in *first*.

Finally, we also add LCP samples to FastLocate. Call *LCP* ((*i, j*)) a function that returns the longest common prefix of the suffix (*i, j*) with the suffix above it (*ϕ*((*i, j*))). In order to compute this function, we add *sampleslcp* to FastLocate, where *sampleslcp*[*i*] is the LCP sample of the suffix at the top of the *i*-th run in the GBWT. Then, *LCP* ((*i, j*)) = *sampleslcp*[*first*[*j* + *i*(*lmax*)].*runID*] − (*j* −*j*), where (*i, j*) is the predecessor of (*i, j*) in *first*.

These additional capabilities are only used in the new set maximal match query algorithm in the last section, when reporting found set maximal matches. See Algorithm 6 for pseudocode of match outputting given an FL and FLCP. The *LCP* and *ϕ* functions remove the need to perform find and extract queries on the GBWT. As before, we need to keep track of the suffix array value at *ak* for all *k*, call this value *sk*. Now, when the length of the maximum LCP of suffix *i* of the reversed path is greater than or equal to the length of the maximum LCP of suffix *i*− 1, we output the matches starting at *i* by iterating up and down the GBWT using the *ϕ* and *ϕ*^−1^ functions. Using the *LCP* and the minimum LCP found in the current direction so far, we obtain the LCP of each suffix with *Q*^∗^ and output it if it is equal to the maximal LCP value. The find queries that FLCP removes the need for used to take *O*(|*Q*| ^2^*tLF*) time in total. Furthermore, the LCP samples remove the need for the query to iterate through the endmarker runs to compute the length of a path in the GBWT. Therefore the time complexity for this version of the algorithm is 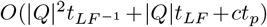, where 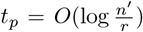 in the GBWT implementation. The suffix array samples at the top of runs and the LCP samples can each be stored in a predecessor data structure with *r* elements and a universe of size *n*. Therefore they take total size *O*(*r*) and answer queries in 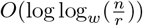 time. This results in a query time of 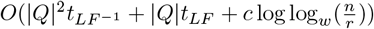.

##### Algorithm 6

OutputMatches: Output Matches using a GBWT, FastLocate, and FastLCP given virtual insertion positions *a*, suffix array values *s*, and LCP values *lcpa, lcpb*

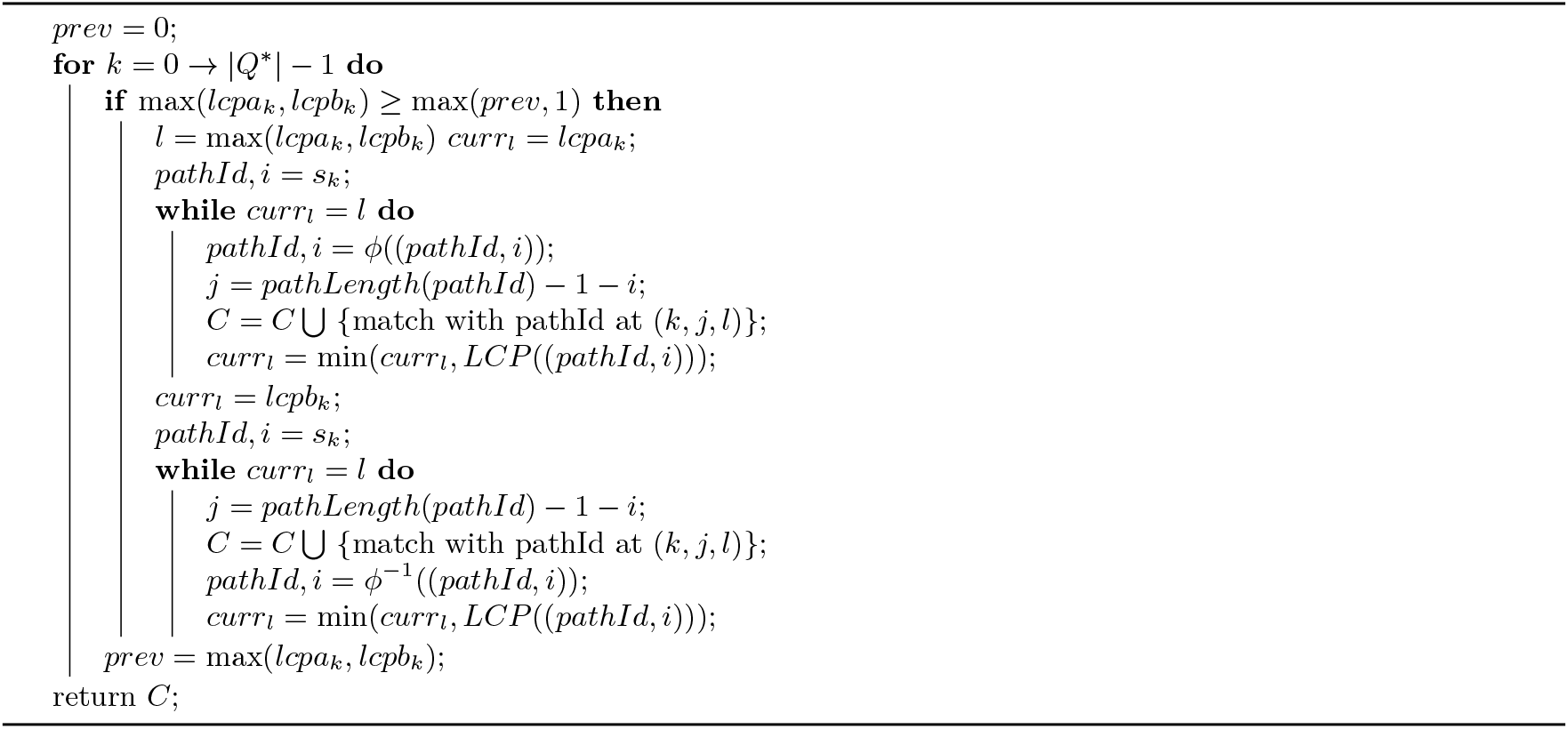

#### FLF + FL + FLCP: Replacing GBWT with Fast LF

The fourth version of the set maximal match query algorithm modifies the GBWT directly. The GBWT currently stores the runs directly and computes LF in a record, *BWT*_*v*_, by iterating through the runs in linear time. Here we add data structures to the GBWT to allow the computation of the LF function with better time complexity. The primary capabilities of the GBWT that the set maximal match query algorithm uses are the *LF* ^−1^ and the generalized *LF* functions. The generalized *LF* function, *LF* ((*v, i*), *u*) for *u* ∈ *Σ*_*v*_ can be computed with the ability to compute rank on *BWT*_*v*_. Then, *LF* ((*v, i*), *u*) = (*u, BWT*.*rank*(*v, u*) + *rank*_*u*_(*BWT*_*v*_, *i*)), where *ranku*(*BWT*_*v*_, *i*) is the number of occurrences of *u* in *BWT*_*v*_[0, *i* − 1]. The natural data structure for this problem is the wavelet tree. The wavelet tree was introduced by Grossi et al. [35]. It is a data structure that allows the computation of *access*(*i*), *rank*_*q*_(*i*), and *select*_*q*_(*i*) in *O*(log *σ*) time for a sequence, *S* = *s*_0_, *s*_1_, *s*_2_, …, *s*_*n*−1_, with an alphabet size of *σ* while using *O*(*n* log *σ*) bits of space [35, 51]. *access*(*i*) = *S*[*i*], *rank*(*i*) is the number of occurrences of *q* in *S*[0, *i*− 1], and *select*_*q*_(*i*) is the position of the *i*-th occurrences of *q* in *S*. Therefore, the generalized *LF* function, would be able to be directly computed for *u*∈ *Σ*_*v*_ if a wavelet tree of *BWT*_*v*_ was stored. However, this would increase the space complexity of the GBWT from *O*(*r*) to *O*(*n*), where *n* is the sum of the lengths of *BWT*_*v*_ for all *v* (the length of the text). Therefore, we instead store a wavelet tree on a sequence representing the runs in *BWT*_*v*_. The wavelet tree is built on the sequence *S* of length *rv*, the number of runs in *BWTv. S*[*i*] = *BWT*_*v*_.*runs*[*i*].*first*, the value of the *i*-th run in *BWTΣ*. Furthermore, for each pair (*u, i*), representing the *i*-th run of *u* in *BWT*_*v*_, the number of occurrences of *u* before this run is stored. Lastly, a data structure is stored to allow the computation of *run*(*BWT*_*v*_, *i*), the index of the run in *BWT*_*v*_ that contains position *i*. This can be done with a predecessor data structure on the starts of the runs of *BWT*_*v*_. Now, to compute *LF* ((*v, i*), *u*) for *u* ∈ Σ, first *run*(*BWT*_*v*_, *i*) is computed, then the number of runs of *u* before *i* can be computed with *rankΣ* (*u*)(*run*(*BWT*_*v*_, *i*)) using the wavelet tree stored at this node. This value along with the *BWT*.*rank*(*v, u*) value can be used to precisely compute *LF* ((*v, i*), *u*) after determining if the value at (*v, i*) is *u* (using the wavelet tree).

A wavelet tree is not the only method of computing rank on *BWT*_*v*_ for *u* ∈ *Σ*_*v*_. An alternative is a small modification of the structure described in Theorem 2.1 by Gagie et al. [30]. Gagie et al. described the RLFM+ data structures by Mäkinen et al. [44] for computing LF using Belazzougui and Navarro’s [7] predecessor data structure. Call the *L*_*GBW T*_ array the *L* array after applying *Σ*_*v*_ to every value. This is equivalent to the concatenation of *BWT*_*Σ*_ for all *v ∈ V*_*Σ*_. The *L*_*GBW T*_ array has alphabet size 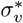, where 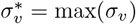. Then, each node *v* must have an associated predecessor data structure with universe size *V*_*Σ*_ and *σ*_*v*_ elements, this structure computes the *Σ*_*v*_ function (whether a node-character is in the local alphabet of *v* and its mapped value). Call the total sizes of the local alphabets *tl* = *Σ*_*v*∈*V*Σ_ *σ*_*v*_. If the universes are concatenated into one predecessor structure, it takes *O*(*tl*) space and 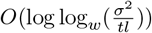 time to compute the *Σ*_*v*_ function (note, *Σ* ≤ *tl* ≤ *r*). Given these structures, the generalized LF function can be computed in 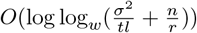. This structure has the same complexity as that of Gagie et al. but may be smaller in practice due to the reduction of the alphabet. Computation of inverse LF takes 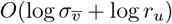 where *v* is the node the function is called from, *u* is the result.

This modification of the GBWT doesn’t directly change the set maximal match query algorithms, it just changes the calls to the LF and inverse LF functions. Therefore, the time complexity of the set maximal match query algorithm remains 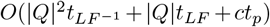 However, with the wavelet tree, *t*_*LF −*1_ has gone from 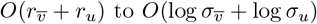 to 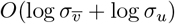 and *t*_*LF*_ has gone from *O*(*r*_*v*_) to *O*(log *Σ*_*v*_ + *t*_*p*_), where *v* is the node the function is called from, *u* is the result, and *t*_*p*_ is the time needed for computation of a predecessor query (to determine if a node is in the local alphabet). With the structure described by Gagie et al., *tLF*^−1^ is now 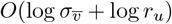 and *t*_*LF*_ is 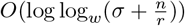.

### FLF + FL + FLCP + CT: Adding the Compressed Text

The block tree is a data structure introduced by Belazzougi et al. that provides access to a text in time 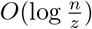 in 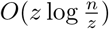 space, where *n* is the length of the text and *z* is the number of phrases the text is parsed into by the Lempel-Ziv algorithm [6]. In their r-index paper, Gagie et al. describe a variant of block trees that allows 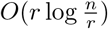 time access to the text while using 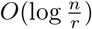 space, where *r* is the number of runs in the BWT of the text [30]. In this version of the set maximal match algorithm, we add this variant of block trees to the GBWT to allow set maximal match query with small space and time complexity bounds.

#### Algorithm 7

LCP: Compute LCPs given a compressed text data structure *CT*, a FastLocate structure, a FLCP structure, add Virtual Insertion positions*a* of string *t* and their corresponding suffix values *s*.

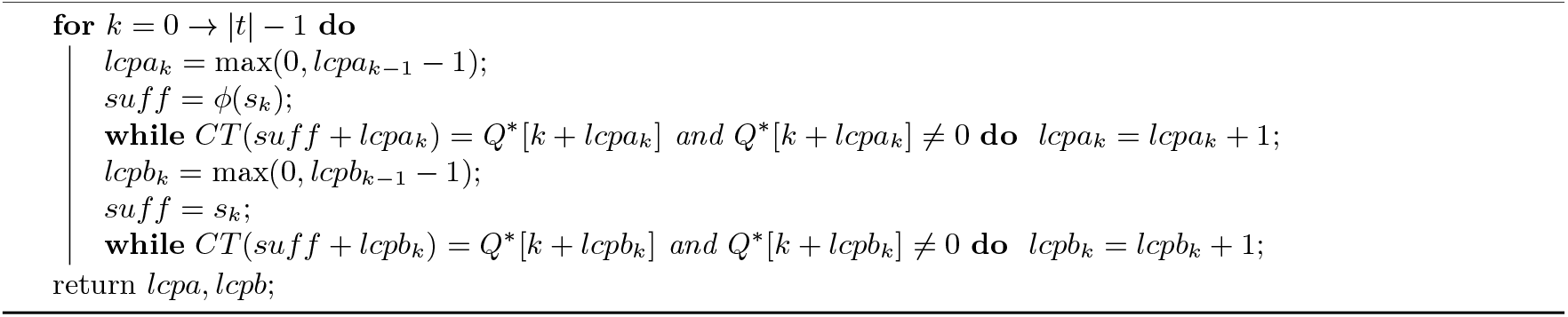

Direct access to the text is used in many set maximal match query algorithms to compute the LCP values efficiently [5, 18, 58, 61]. The key property is that if suffix *i* of a query string *s* has an LCP of *j* with the suffixes of the text, then the suffix *i* + 1 of the query string will have an LCP of at least *j* − 1. Therefore, while computing the LCP values for suffix values *i* = 0 → |*s*| − 1, a linear scan down *s* is all that is needed. An analogous property holds for query paths and the LCPs of their reversed prefixes with the GBWT. Therefore, we use the block tree variant of Gagie et al. to access the text. A linear amount of text accesses results in 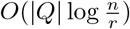 time complexity for computing all the LCPs given the virtual insertion positions. See Algorithm 7 for pseudocode of computation of LCPs with linear text accesses.

This final modification of the GBWT results in a set maximal match query time of *O*(|*Q*| (*t*_*acc*_ +*t*_*LF*_)+*ct*_*p*_) where 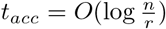 is the time to access one value of the text. Note however that this modification is the only one to increase the space complexity of the GBWT. This modification increases the space complexity of the GBWT from *O*(*r*) words to 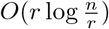. If the data structures described previously are used, the time complexity is 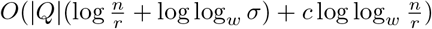.

Note that our implementation is slightly modified from the original description by Gagie et al. We store four non-overlapping half blocks for every sampled location at every level instead of seven (some overlapping) half blocks. This is designed to support point queries in 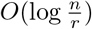 time while reducing the size of the compressed text data structure. We also prune the sampled locations at every level, only keeping those mapped to from the previous level. Furthermore, all half-blocks with equal underlying text values are mapped to the same location in the next level. Finally, note that a possible way to improve the size of the structure is to choose the set of sampled locations at the next level as a minimal set cover of the half blocks of the previous level (greedy minimal set cover might be practical here). We did not implement the last optimization.

### 3.3 Long Match Query Algorithm in GBWT

Here, we provide an algorithm that outputs all long matches between a query path-string and path-strings in a GBWT. We first solve the problem where a match, (*i, j, k*), between a query path-string *Q* and a path-string *sp* ∈ *S*_*P*_ is a long match if it is locally maximal and *k* ≥ *L*. We then show that this algorithm can be easily extended to the case where the weight of a string is nonincreasing over the suffixes of *Q*. I.e. for any weight function, 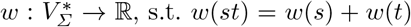 and *w*(*s*) ≥ 0 for all strings *s, t*. This applies to the pangenome graph long match query, where long matches are being searched for on the base-strings (each node has a sequence associated with it and the weight of each node is the length of the sequence).

Here, we first describe the algorithm where a locally maximal match is a long match if the length of the number of node-characters in the match is at least *L*. The algorithm operates in three high level steps. First, “virtual insertion” positions are computed for each suffix of the reversed query. Then the length of the longest common prefix with all suffixes in the GBWT of every suffix of the reversed query is computed. Finally, the long matches are found and outputted using these computed values. We refer to *Q*^∗^ as the endmarker appended to the reverse of *Q*, (*Q*^∗^ = rev(*Q*)0).

The *k*-th suffix of |*Q*^∗^| is *Q*^∗^[*k*.. |*Q*^∗^|− 1]. We refer to the length of the longest common prefix of the *k*-th suffix of *Q*^∗^ with the rest of the suffixes in the GBWT as 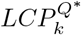. Then, the “virtual insertion” position of the *k*-th suffix is any position (*v, i*) such that the suffix at (*v, i*) or (*v, i* −1) has a longest common prefix with the *k*-th suffix of *Q*^∗^ with length 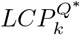. We refer to the virtual insertion position of suffix *i* of *Q*^∗^ as *a*_*k*_. Therefore, if the suffix range (as defined in [38]) in the GBWT of the prefix of length 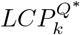 of the *k*-th suffix of 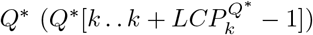 is [(*v, l*), (*v, r*)], then *a*_*k*_ is any value in [(*v, l*), (*v, r* + 1)].

Note that using certain virtual insertion values to compute matching statistics can allow the computation of matching statistics (and therefore set maximal matches and MEMs) without thresholds (introduced by Bannai et al. [5] and construction shown by Rossi et al. [58]). See Section 3.2 for further discussion. The computation of virtual insertion position and LCP lengths are exactly the same as in the set maximal match query algorithm, and are discussed in Section 3.2. Therefore, the first two steps of the long match query algorithm can be completed by performing exactly the set maximal match query algorithm except for the outputting of set maximal matches. The time complexity of obtaining the virtual insertion positions and the lcp values is the same as that of the set maximal match query algorithm version used to obtain the values without the term containing *c*.

The third step of the long match query algorithm has a basic high level idea: do a sweep from suffix |*Q*^∗^| 1 to 0 of *Q*^∗^ in the GBWT. During the sweep, given the block of suffixes in *BWT*_*Q∗* [*i*]_ that have a common prefix with suffix *i* of *Q*^∗^ length *L* or longer, compute the block of suffixes in *BWT*_*Q∗* [*i* − 1]_ that have a common prefix with suffix *i* − 1 of *Q*^∗^ length *L* or longer and output long matches corresponding to all suffixes *j* of the text that are contained in the block of *BWT*_*Q∗* [*i*]_ while suffix *j* − 1 of the text is not contained in the block *BWT*_*Q∗* [*i* −1]_. If this is possible, then all long matches can be outputted by iterating from suffix *i* = |*Q*^∗^| → 1 0 maintaining these blocks and outputting matches with the suffixes that leave them. We denote the block of suffixes in the GBWT that match with suffix *i* of *Q*^∗^ length *l* or more as 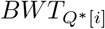 is the first suffix in the block and 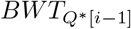 is the first suffix after the start of the block not in the block. Note that the suffix array is not actually stored and *SA*[(*v, i*)] represents the suffix that would be in the suffix array at position *C*[*v*] + *i*. Then, given 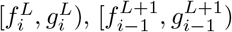 can be computed directly from the LF function, 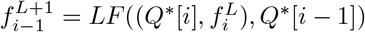 and 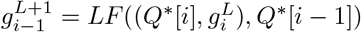. If the block is empty at this point, use the previously computed virtual insertion positions and LCP lengths to initialize it if needed. Finally, we iteratively add adjacent suffixes that have a common prefix with *Q*^∗^[*i* − 1, |*Q*^∗^| − 1] of length *L* to the block until no more such adjacent suffixes exist. At that point, 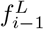 and 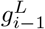 have been computed. If the GBWT has the LCP samples data structure, this process takes 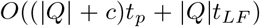 time over the whole long match query, where *c* is the number of matches outputted.

The only thing left for the complete long match query is the outputting of matches with suffixes that have left the block from one suffix to the next. For two blocks, 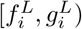 and 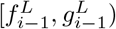, for any suffix 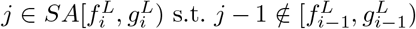, (say suffix *j* of the text corresponds to suffix *k* of the reversed path-string *sp* in the text) then *Q*^∗^ and *sp* have a long match equal to (*i*; *k*;*LCP*(*Q*\*[*i*; |*Q*\*|−1]; *sp*[*k*; |*sp*| − 1])).. All such suffixes are located at exactly the positions 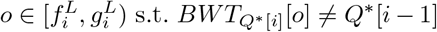. Therefore, all the suffixes that leave the block 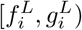 can be found by iterating through the runs in the block in *BWT*_*Q∗* [*i*]_ and outputting all the long matches corresponding to the suffixes in runs with values not equal to *Q*^∗^[*i* −1]. However, note that the length of these matches is not trivial to compute efficiently. This is addressed by maintaining a dynamic predecessor data structure on the suffixes within the block. Whenever suffix *k* of the text is added to the block at suffix *p* of *Q*^∗^, then the value *k* −*p* is inserted into the predecessor data structure, while *p* + *L* is associated with this value in the data structure as the start position in *Q*^∗^ of this match. The value of *k* − *p* remains constant while this suffix and the query path-string match. Therefore, while computing the long matches to output at some block 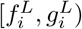, if 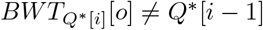, then if the start position associated with the value *SA*[(*Q*^∗^[*i*], *o*)]− *i* is *s*, the length of the match to be outputted is *s*−*i*. We also remove the value *SA*[(*Q*^∗^[*i*], *o*)]−*i* and its associated start position from the dynamic predecessor data structure when outputting matches to *SA*[(*Q*^∗^[*i*], *o*)]. The outputting of the matches takes time equal to the time needed for iterating through the runs in the block and for recovering the suffix array values of the outputted matches. If the GBWT has the suffix array samples as described in Section 3.2 and random access to the runs, then these time complexities are *O*(*c*) and *O*(*ct*_*p*_) respectively over the whole long match query. If the runs are replaced with the predecessor data structures as described in Section 3.2, these time complexities are *O*(*t*_*p*_ |*Q*| + *c*) and *O*(*ct*_*p*_) respectively. Finally the maintenance of the dynamic predecessor data structure takes *O*(*ct*_*dp*_) over the whole long match query where *t*_*dp*_ is the time needed for insertion, deletion, and query in the structure. The dynamic predecessor data structure has a universe of size *n* + *Q*^∗^ and at most *c* elements. Therefore, if exponential search trees by Andersson and Thorup are used, then the dynamic predecessor data structure takes *O*(*c*) space and 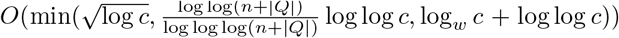 time for query and update [3].

The long match query algorithm provided here has three main stages: virtual insertion position computation, LCP computation, and match outputting. The first two stages are covered in the set maximal match queries. The third operation takes a constant number of *LF* operations per node-character in the query. Furthermore, there is a constant number of *LCP* operations for every match and for every node-character in the query. Finally, for every long match outputted, a predecessor query is inserted to, queried from, and removed from once. Therefore, the time complexity of the third stage of the long match query algorithm is *O*(*Q* (*t*_*LF*_ + *t*_*p*_) + *c*(*t*_*p*_ + *t*_*dp*_)), where *t*_*LF*_ is the time for LF computation, *t*_*p*_ is the time for static predecessor query on *O*(*r*) elements from a universe of size *n*, and *t*_*dp*_ is the time complexity for query and update of a dynamic predecessor data structure with a universe of size *n* and at most *c* elements. *c* is the number of matches outputted, *n* is the size of the GBWT, and *r* is the number of runs in the GBWT. If the virtual insertion and LCP computation from Section 3.2 with the best time complexity is used along with the same predecessor data structures, then the overall time complexity for the long match query described here is 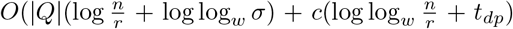 where 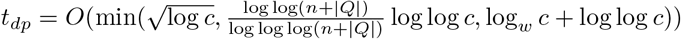.

The long match query algorithms presented in the paper can be understood with Algorithm 8. The only difference in the long match query versions is how the virtual insertions and lcps are computed. The methods of computing virtual insertions and lcps in the long match algorithm will depend on the available data structures. For any set of available data structures, the virtual insertions and lcps are computed the same way that they are computed in the corresponding set maximal match query.

#### Arbitrary Node Weights

Here, we adapt the long match query algorithm to any weight function *w* on strings s.t. *w*(*st*) = *w*(*s*) + *w*(*t*) and *w*(*s*) ≥ 0 for all strings *s, t*. The first step is to compute the value *w*(*Q*^∗^[*i*, |*Q*^∗^| − 1]) for all suffixes *i* of *Q*^∗^. This can be done by a linear amount of calls to the weight function.

##### Algorithm 8

Long Match Query: Outputs all long matches of query path *Q* and a GBWT

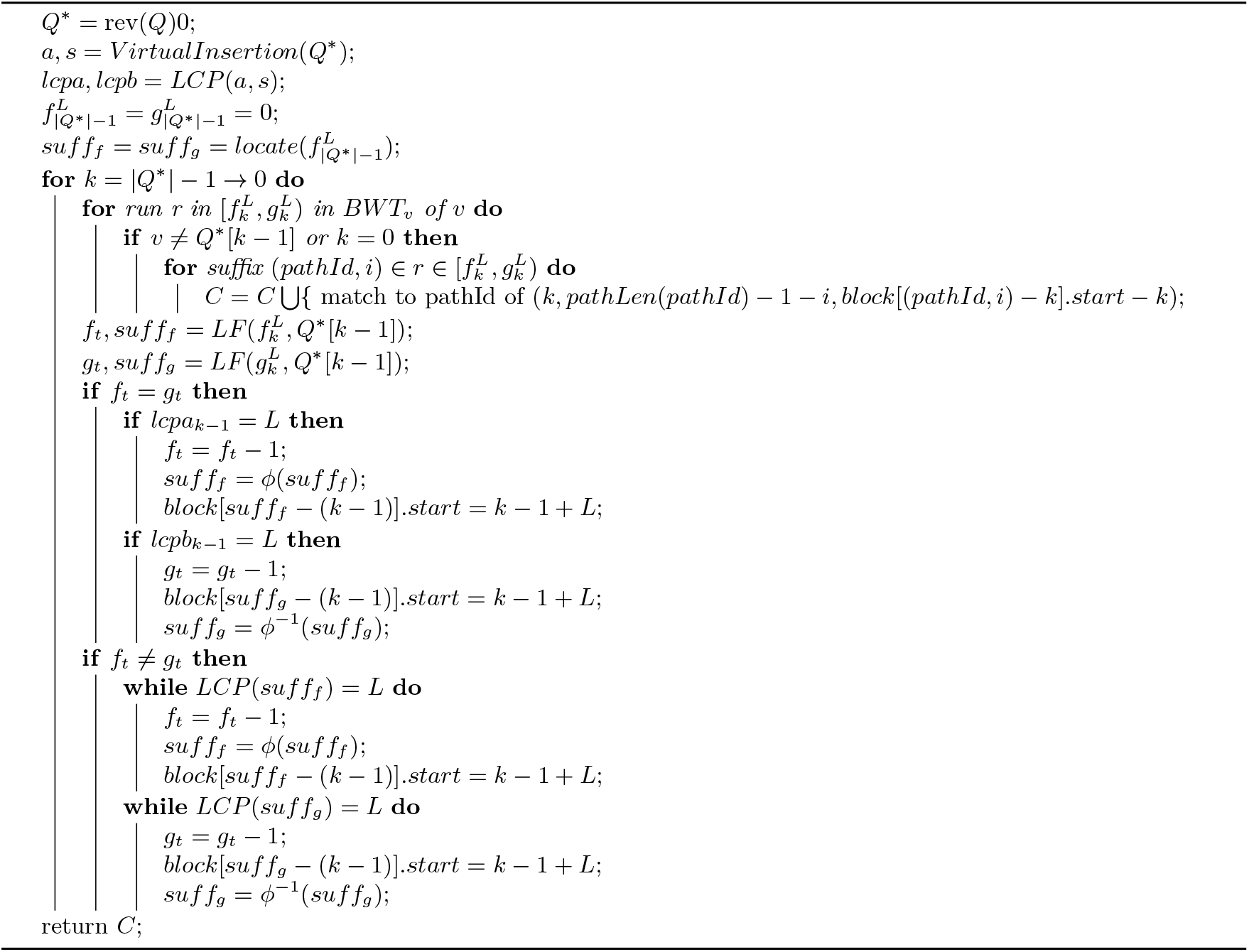

These calls can also be restricted to a single node-character for each call. Then, for each suffix *i*, a value *L*_*i*_ is computed. *L*_*i*_ is the smallest possible value s.t. *w*(*Q*^∗^[*i, i* + *L*_*i*_ − 1])≥*L*. All *L*_*i*_’s can be computed in a linear two point scan of the weights of the suffixes. Note that *L*_*i*_≤*L*_*i*+1_ + 1. Now, only a few changes need to be made to the long match query algorithm. First, instead of computing the block 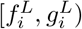 for every suffix *i*, we compute the block 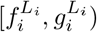 for every suffix *i*. This should be done by iterating from *l* = *L*_*i*+1_ + 1 → *L*_*i*_ iteratively adding the suffixes that have a common prefix of length *l* with the *i*-th suffix of *Q*^∗^. Finally, when a suffix *k* of the text is added to the block, the value added to the dynamic predecessor data structure should be *k* − *i* and its associated start position in *Q*^∗^ should be *i* + *l*. This algorithm has the exact same time complexities as the long match query algorithm with an added *O*(|*Q*|) calls to the weight function.

## 4 Results

We construct the indexes and run the algorithms described in this paper. The implementation is available at https://github.com/ucfcbb/GBWT-Query. We compare the space and time usage of the various versions of set maximal match and long match query described in this paper for GBWTs built on Chromosome 21 of the 1000 Genomes (1000G) project phase 3 release dataset [1]. This dataset has 5008 haplotypes (2504 individuals) and roughly 1.1 million variants. First, a variation graph (vg) was built for the dataset. Then the vg was used to build an xg [32]. Finally, a bidirectional GBWT of this dataset was built using the xg with the vg gbwt sub command and the --discard-overlaps option [32]. This process is as outlined in https://github.com/vgteam/vg/wiki/Index-Construction. This resulted in a GBWT with 10,016 sequences, (a forward and reverse orientation for each haplotype) and a graph with roughly 6.6 million nodes. It had about 26.6 million runs and the sum of lengths of the sequences in it was roughly 26 billion. This demonstrates that the low average outdegree and runs assumption of the GBWT holds in the 1000G dataset. The average runs per node was 4.03 and the average outdegree was 1.22.

Given the bidirectional GBWT of Chromosome 21 of the 1000G dataset, we construct the indexes used for evaluating the space and time usage of the various set maximal match and long match query versions. In order to do this, one hundred haplotypes are selected at random in the GBWT with uniform probability for removal. The forward and reverse orientations of these haplotypes are removed from the GBWT and stored for later query. This results in 200 sequences being removed from the GBWT. Then, the auxiliary data structures were built on this modified GBWT. First the FastLocate (FL) structure, that stores suffix array samples at the top of runs in the GBWT, was built. This structure was implemented by Siren in the GBWT github (https://github.com/jltsiren/gbwt/commit/180e041a029af05f1d74e57058e3c16974c85362) after the publication of the GBWT paper. Then, FastLCP (FLCP), the addition to FastLocate that stores LCP array samples at the top of runs in the GBWT and adds suffix array samples at the bottom of runs is built. FLCP is designed as the corresponding capabilities in the BWT were originally described in the r-index paper [30]. Then, as described in Section 3.2, Fast LF (FLF) is built, a modification of the GBWT where the direct storage of the runs is replaced with storage of a structure that allows efficient computation of rank. Finally, the compressed text (CT) data structure described by Gagie et al. in their r-index paper is built. The implementations of the GBWT and FL are by Siren et al. and the implementations of FLCP, FLF, and CT are by the authors of this paper. Note that in this implementation of FLF, predecessor structures are used, but there are alternatives with the same space and time complexities (for example, wavelet trees or wavelet matrices). These implementations make use of the vgteam’s fork of the SDSL library [32, 34]. Note that we refer to the sampled document array described in the GBWT paper as DA. The sampled document array is sampled every *s* positions in the BWT, see Section 2.1 for more information. For the 1000G Chromosome 21 dataset, we used *s* = 1, 024.

### 4.1 Indexes

In Table 2, the space taken by each data structure for the dataset with 200 sequences removed is reported. The space complexity of each structure and their actual space used is reported. All data structures have a space complexity of *O*(*r*) words except for DA and CT. These have 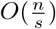 and 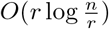 space complexities respectively, where *r* is the number of runs in the *BWT, s* is the document array sample interval, and *n* is length of the text. The FL and FLCP structures are three to four times larger than the GBWT. The FLF data structure is 2.2 times larger and the CT data structure is 8.3 times larger. The CT data structure can be expected to be on the order of 10 times the size of the GBWT since 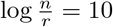 in this case.

**Table 2.** Space Complexity in words of Data Structures: theoretical space complexity and used space for data structures built on a bidirectional GBWT of 1000G Chromosome 21 with 200 sequences removed (100 haplotypes). Refer to Table 1 for more details about these data structures.

|  | DA | GBWT | FL | FLCP | FLF | CT |
| --- | --- | --- | --- | --- | --- | --- |
| Space Complexity | $O(\frac{n}{s})$ | $O(r)$ | $O(r)$ | $O(r)$ | $O(r)$ | $O(r \log \frac{n}{r})$ |
| Space Used (MB) | 71.7 | 84.2 | 248.8 | 292.2 | 184.1 | 698.7 |

### 4.2 Queries

Given the structures built in the previous section, we perform a query benchmark for the versions of set maximal match and long match presented in this paper. For every sequence removed from the original Chromosome 21, each set maximal match query version is performed from the first presented in the paper to the last. Then long match query is performed with varying supporting data structures. Each long match query computes virtual insertions and their corresponding LCPs the same way the set maximal match query that uses the same data structures does. Note that the set maximal match and long match queries also have another version tested. This one computes LCP values using access to the text through CT, but computes LF using the GBWT. This is essentially the final version of each algorithm with LF computed by the GBWT instead of FLF. Furthermore, all empty nodes in the GBWT are excluded from the query path.

For each query performed, the matches and the time taken were measured. In Table 3, the average time and standard deviation of the 200 queries can be seen for each algorithm run. The long match query results are for *L* = 7, 500. The space needed for the supporting structures is also provided. We first discuss the results on set maximal match queries. The first thing to note is between the first and second set maximal match query version results. While GBWT+FL has a theoretically better time complexity, the difference is only in changing a *cs* term to 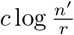 (see Section 2.2). This is not the dominating factor in the time complexity, the dominating factor is the 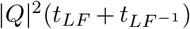. Also note that when *s* is chosen to be 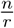, as it is in this case, the space complexity of GBWT+DA remains *O*(*r*) with samples. The FLCP structure removes the need to perform *Q* ^2^ LF calls, speeding up the query with a little more space cost. The purpose of the FLF data structure was to reduce the complexity of computing the *LF* function. In practice, it turns out the GBWT computes LF faster due to the very low number of runs per node. The fifth set maximal match query adds access to the text through CT and implements the final set maximal match query version presented in this paper. The removal of the *Q* ^2^ inverse LF computations fails to compensate for the slowness of computation of LF using FLF. The final set maximal match query version uses the benefits of CT and GBWT. It is the same as the previous algorithm but LF is computed by the GBWT. This is the fastest by far. Also note that while the CT data structure is large, it removes the bidirectional requirement for the GBWT. Therefore, the space requirements for the last two versions can be roughly halved if a non-bidirectional GBWT is used (and its corresponding structures).

**Table 3.** Set Maximal and Long Match query times in seconds. The total space for the required structures is also provided. The first five rows of the set maximal match query correspond to the five versions presented in Section 3.2 in order. The only difference in the long match query versions are the computation of virtual insertion positions and their corresponding LCP values. This is done in the same fashion as in the set maximal match query that utilizes the same data structures. Long match queries are run with *L* = 7, 500. The last row is the second last row with the LF function computed by the GBWT instead of FLF.

| Data Structures | Space (MB) | Set Max Match (s) |  | Long Match (s) |  |
| --- | --- | --- | --- | --- | --- |
|  |  | Mean | Std. Dev. | Mean | Std. Dev. |
| GBWT+DA | 156.0 | 10.27 | 1.20 |  |  |
| GBWT+FL | 333.0 | 9.56 | 1.15 |  |  |
| GBWT+FL+FLCP | 625.2 | 8.32 | 1.00 | 9.65 | 2.58 |
| FLF+FL+FLCP | 725.1 | 164.66 | 43.82 | 215.29 | 66.06 |
| FLF+FL+FLCP+CT | 1423.8 | 70.58 | 16.08 | 130.30 | 41.31 |
| GBWT+FL+FLCP+CT | 1323.9 | 3.66 | 0.85 | 5.63 | 1.90 |

In Table 4, the average and standard deviation of the number of matches is reported for the set maximal and long match query algorithms. Results for various values of *L* are provided for the long match query. Note that in the GBWT of Chromosome 21 of 1000G, the first roughly 750,000 nodes in every forward oriented path are equal (and therefore the last roughly 750,000 nodes in every backward oriented path are equal). As can be seen in Table 4, each long match query for *L* = 200, 000 returned exactly 4,908 matches. These are all matches at the beginning of the chromosome with every path in the GBWT that is oriented in the same direction as the query path. The average run time for varying *L* is plotted for each version of the long match query in Figure 3 along with the average number of matches outputted. The relative speed of the queries with varying combinations of data structures remains consistent over varying L and between long match and set maximal match queries. In general, the long match queries slow down as *L* gets smaller due to a larger number of outputted matches.

**Table 4.**
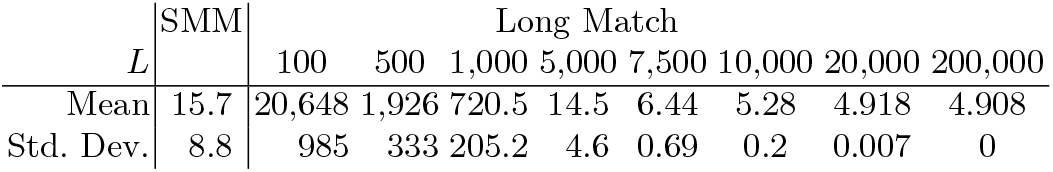
Mean and Std. Dev. of # Matches in thousands for set maximal match query and long match query for various values of *L*. SMM represents set maximal query. The results are for 100 randomly selected haplotypes out of 5008 from the 1000G dataset on Chromosome 21. Queries were performed for both orientations of haplotypes in a bidirectional GBWT.

| $L$ | SMM | Long Match | | | | | | | |
| --- | --- | --- | --- | --- | --- | --- | --- | --- | --- |
|  |  | 100 | 500 | 1,000 | 5,000 | 7,500 | 10,000 | 20,000 | 200,000 |
| Mean | 15.7 | 20,648 | 1,926 | 720.5 | 14.5 | 6.44 | 5.28 | 4.918 | 4.908 |
| Std. Dev. | 8.8 | 985 | 333 | 205.2 | 4.6 | 0.69 | 0.2 | 0.007 | 0 |

**Fig. 3.**
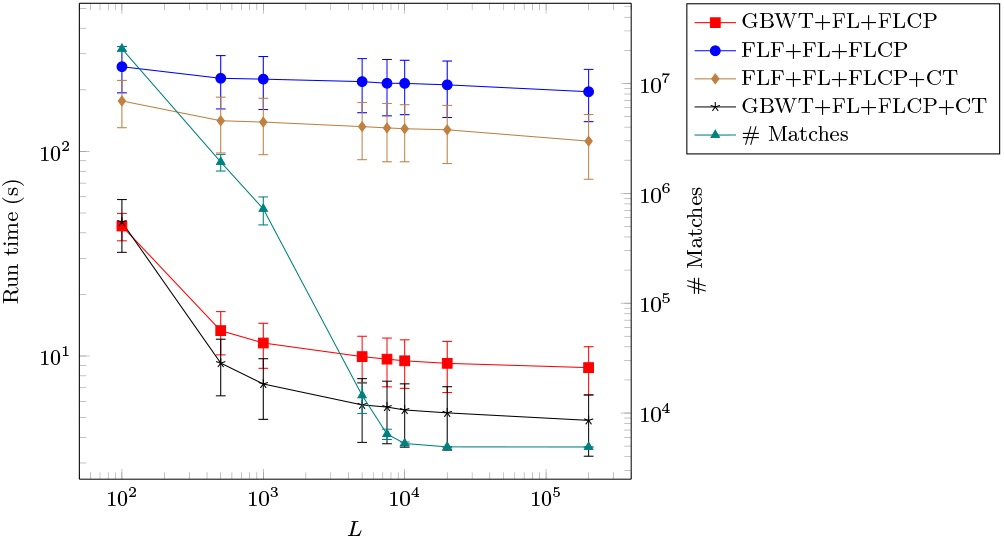
Average run times for versions of the long match query with varying *L*. Error bars are one standard deviation from the mean. The average number of matches per query is plotted as well.

## 5 Discussion

In this paper, we have described set maximal match and long match query algorithms on the GBWT. Five versions of set maximal match query have been provided to allow the computation of set maximal match query in scenarios where certain data structures are not available or memory is a constraint. By relaxing the constraints for the computed matching statistics, the set maximal match queries provided here don’t require the thresholds previous set maximal match queries on the BWT require [5]. Various long match query algorithms for different combinations of available data structures are provided as well. We provide these long match and set maximal match query algorithms on the GBWT through the application of the r-index data structures to the GBWT. Extending these algorithms to use new combinations of the data structures described here is straightforward. The most efficient set maximal match and long match queries presented here require 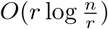 space for the supporting structures and run in 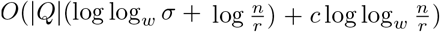 and 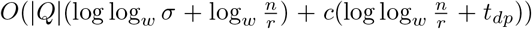 time respectively, where 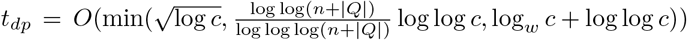.

Given the similarities between the GBWT and BWT, the queries presented here applies to the BWT as well. In particular, the long match query on the BWT is novel. The long match query algorithm presented here is the first efficient long match query algorithm on the BWT the authors are aware of. Possible directions for future work include speeding up the FLF data structure or reducing the space needed for the compressed text data structure. The size of the compressed text data structure can be reduced by pruning of the tree. This may be done by selecting a small set of nodes at every level such that every node at the previous level can be mapped from. This set can be chosen using greedy set cover. Another possible direction is making the structures described here dynamic. This should be straightforward using dynamic predecessor data structures and techniques similar to the dynamic Burrows-Wheeler transform and d-PBWT [60, 61]. The ordering of the haplotype paths in the text may be evaluated as well. A modification of optBWT may be able to output the optimal ordering of haplotype paths in the text for the least amount of runs in the GBWT in linear time [8, 15]. Another possibility is making the GBWT independent of haplotype path order in the text in a similar fashion to the extended r-index [11, 45]. The optBWT approach may result in fewer runs [16].

Finally, in this paper we defined relationships between maximal match types in the PBWT and BWT paradigms. Using these relationships, we defined maximal matches in the GBWT and described their similarities to matches in the BWT and PBWT. We also defined new maximal match types in the BWT paradigm analogous to some PBWT match types. Efficient algorithms for outputting TEMs, long TEMs, or only outputting matches that are both TEMs and MEMs are possible directions for future work.

## Availability

The source code for GBWT-Query is available at https://github.com/ucfcbb/GBWT-Query.

## Acknowledgments

This work was supported by the National Institutes of Health under award number R01HG010086.

## Footnotes

1 Here, generalized FM-Index refers to an FM-Index for a set of strings. It is an FM-Index built on the concatenation of the strings with an endmarker appended to every string. The GBWT is referred to as an emulated FM-Index because while the same operations are supported, the BWT is not move to front encoded and the data structures that support computation of *BWT*.*rank*(*v, w*) + *BWT*.*rank*((*v, i*), *w*) (equivalent to *Occ*(*w*, (*v, i −* 1) as defined in [26]) are different in the GBWT and FM-Index, resulting in different time and space complexities.

